# Rhizobial effector NopM ubiquitinates Nod factor receptor NFR5 and promotes rhizobial infection in *Lotus japonicus*

**DOI:** 10.1101/2024.04.28.591553

**Authors:** Yanan Wang, Hanbin Bao, Yutao Lei, Lifa Yuan, Haoxing Li, Hui Zhu, Dawei Xin, Christian Staehelin, Yangrong Cao

## Abstract

Bacterial pathogens and nitrogen-fixing rhizobia employ type III protein secretion system (T3SS) effectors as potent tools to manipulate plant signaling pathways, thereby facilitating infection and host colonization. However, the molecular mechanisms by which rhizobial effectors affect legume infection remain largely elusive. In this study, we investigated the symbiotic role of T3SS effectors in the interaction between *Sinorhizobium fredii* NGR234 and *Lotus japonicus*. Mutants deficient in the T3SS genes *Tts1* or *NopA* showed enhanced rhizobial infection of *L. japonicus* roots. Further mutant analysis showed that the NopT effector negatively affects infection, while the NopM effector in the absence of NopT promotes infection. Notably, NopM interacts with the Nod factor receptors LjNFR1 and LjNFR5. NopM ubiquitinates LjNFR5 on ten lysine residues as identified using mass spectrometry. Expression of *NopM* in *L. japonicus* resulted in an approximately twofold increase in LjNFR5 protein levels and enhanced rhizobial infection. Our findings indicate that NopM directly interferes with the symbiotic signaling pathway through interaction and ubiquitination of Nod factor receptors, which likely benefits rhizobial infection of *L. japonicus*. Our research contributes to the intricate interplay between the Nod factor signaling pathway and rhizobial T3SS effectors delivered into host cells.

**Significance Statement:** This study unravels intricate mechanisms underlying the symbiotic interaction between *Sinorhizobium fredii* NGR234 and *Lotus japonicus*. By focusing on the role of the rhizobial effector NopM, this research sheds light on how bacteria manipulate host symbiotic signaling to promote infection and colonization. NopM interacts with NFRs and ubiquitinates LjNFR5, suggesting a novel strategy employed by rhizobia to enhance symbiotic signaling.

## Introduction

The legume-rhizobium symbiosis leads to the formation of nodules, where rhizobia reside and reduce nitrogen into ammonia for use by the plant host. Nodule formation depends on complicated regulations and mutual signal exchanges between hosts and symbionts. Legume roots release flavonoids, which activate NodD transcriptional regulators, initiating the expression of genes for type III protein secretion system (T3SS) and biosynthesis of lipo-chitooligosaccharides known as Nod factors (NFs) (1). These NFs are recognized by Nod factor receptors (NFRs) in host plants, such as LjNFR1 and LjNFR5 in *Lotus japonicus*, activating symbiotic signaling pathways crucial for successful nodulation (2).

Similar to gram-negative bacterial pathogens whose type III protein secretion system (T3SS) effectors are vital for interacting with host pathways, rhizobia inject effector proteins into plant cells to promote rhizobial symbiosis and/or suppress plant immunity (3). The expression of rhizobial genes related to T3SS and effectors is induced by flavonoid-NodD complex and and the positive regulator TtsI (4). These T3SS components and effectors, termed nodulation outer proteins (Nops), have been implicated in infection, nodulation, and host specificity (3, 5). The symbiosis promoting effects of translocated effectors are often attributed to the suppression of host defense reactions, while negative effects may be the result of direct or indirect recognition of effectors by intracellular resistance proteins (3).

Several Nops have been characterized biochemically to date. For instance, *Sinorhizobium fredii* NGR234, a strain with broad-host range, has been well-studied. NopA (6) and NopB (7) are major components of the T3SS complex and interact physically with NopX, facilitating the translocation of effector proteins into plant cells (8). NopL, known to delay nodule senescence in bean, acts as a mitogen-activated protein kinase (MAPK) substrate and reduces levels of pathogenesis-related (PR) proteins in plants (9, 10). NopP synergistically promotes nodulation with NopL and is phosphorylated by unknown kinases (11). NopT, a cysteine protease, regulates symbiosis between NGR234 and various hosts (12), triggering rapid cell death dependent on its protease activity (12). Additionally, NopT cleaves both LjNFR5 and GmPBS1, regulating plant symbiotic signaling and defense responses, respectively (13, 14). NopM, an E3 ubiquitin ligase, dampens flagellin-induced immunity and interferes with MAPK signaling (15).

While effectors produced by bacterial pathogens are known to target plant components of pattern-triggered immunity pathway to favor pathogenic infection, it remains unknown whether rhizobial effectors function similarly regulating symbiotic signaling pathway. Here, we investigated the function of T3SS effectors from *S. fredii* NGR234 in the interaction with *L. japonicus* and found that the T3SS negatively affects rhizobial infection in this plant. Further mutant analysis showed that NopT suppressed rhizobial infection, whereas NopM (in the absence of NopT) promoted infection. NopM interacts with LjNFR1 and LjNFR5. NopM selectively ubiquitinates LjNFR5, increasing LjNFR5 protein levels. Transgenic plants expressing NopM showed increased infection events and a higher number of nodule primordia. These findings establish a direct link between rhizobial effectors and the symbiotic signaling pathway, providing crucial insights into the mechanisms of legume-rhizobium symbiosis.

## Results

### NopM in a Δ*nopT* mutant positively regulates rhizobial infection of *L. japonicus*

To investigate the involvement of effectors secreted by the T3SS in the regulation of rhizobial infection, we generated two T3SS knockout mutants of *S. fredii* NGR234, namely *ΩttsI* and *ΩnopA*. The former lacked the transcriptional regulator gene *TtsI* (*ΩttsI*), and the latter lacked a component of the T3SS pilus (*ΩnopA*). Surprisingly, both mutants induced a significant increase in infections of *L. japonicus* (Gifu) compared to the wild-type strain (Fig. 1A, Fig. 1B and Fig. S3A). These findings suggested that T3SS in NGR234 negatively influences rhizobial infection.

**Figure 1.**
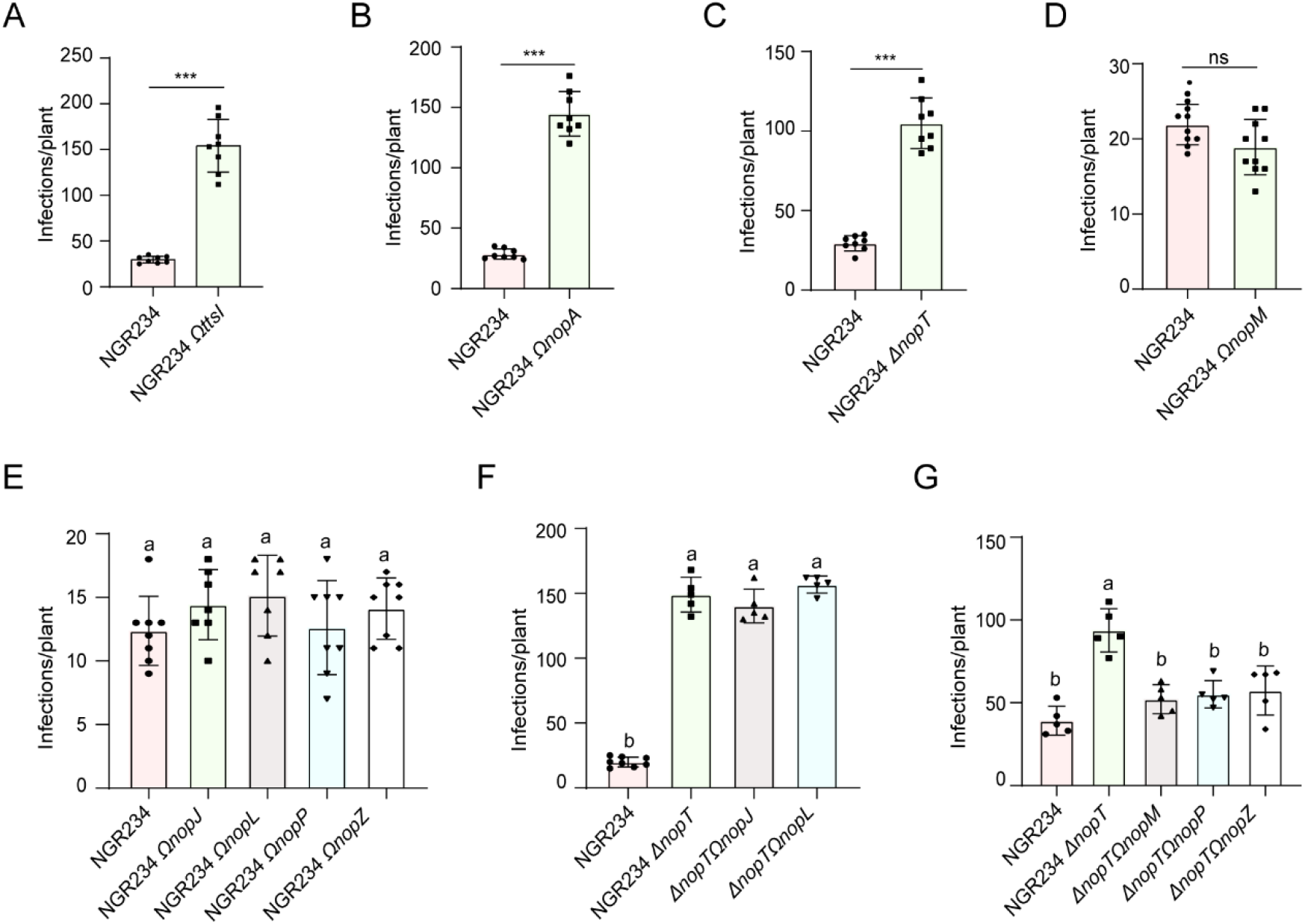
Infection Numbers of *L. japonicus* Roots Inoculated with NGR234 and Mutant Strains. The number of rhizobial infections formed on *L. japonicus* (Gifu) roots inoculated with *S. fredii* NGR234 wild-type and mutants at 6 dpi, (A) NGR234 (n=8) and *ΩttsI* (n=8); (B) NGR234 (n=8) and *ΩnopA* (n=8); (C) NGR234 (n=8) and Δ*nopT* (n=8); (D) NGR234 (n=10) and *ΩnopM* (n=10); (E) NGR234 (n=8), *ΩnopJ* (n=7), *ΩnopL* (n=7), *ΩnopP* (n=8), and *ΩnopZ* (n=8); (F) NGR234 (n=8), Δ*nopT* (n=5), Δ*nopTΩnopJ* (n=5), and Δ*nopTΩnopL* (n=5); (G) NGR234 (n=5), Δ*nopT* (n=5), Δ*nopTΩnopM* (n=5), Δ*nopTΩnopP* (n=5), and Δ*nopTΩnopZ* (n=5). All the rhizobia were labeled with GFP markers. Data are mean ± SD ( Student’s t-test, P < 0.01; ns, not significant).

To explore the specific contribution of individual effectors to rhizobial infection, we examined mutants for six putative or *bona fide* effector genes (*ΩnopJ, ΩnopL, ΩnopP, ΩnopZ, ΩnopM*, Δ*nopT*) (16). Compared to the parent strain NGR234, the Δ*nopT* mutant showed increased numbers of infection (Fig. 1C and Fig. S3B), however, no significant differences were observed for other effector mutants (Fig. 1D, 1E and Fig. S3B). This suggested that the impact of effectors on symbiosis may not be dependent on a single effector but rather on the combined consequences of their activities, acting synergistically, redundantly, or antagonistically.

To identify potential effectors that positively regulate rhizobial infection in the Δ*nopT* mutant, we constructed five double mutants (Δ*nopTΩnopJ*, Δ*nopTΩnopL*, Δ*nopTΩnopP*, Δ*nopTΩnopZ*, Δ*nopTΩnopM*) and inoculated them onto *L. japonicus*. The Δ*nopTΩnopJ* and Δ*nopTΩnopL* double mutants induced massive infections similar to Δ*nopT* (Fig. 1F and Fig. S3C), indicating that NopJ and NopL in the Δ*nopT* mutant might not affect rhizobial infection. In contrast, Δ*nopTΩnopP*, Δ*nopTΩnopZ*, and Δ*nopTΩnopM* double mutants showed reduced infection numbers similar to the NGR234 WT strain (Fig. 1G), suggesting that NopM, NopP and NopZ might act antagonistically with NopT in rhizobial infection. Despite previous reports on NopM being an E3 ligase involved in symbiosis regulation (15, 17), its host targets have not been reported. We then focused on the identification of NopM targets in plants.

### NopM interacts with LjNFR1 and LjNFR5

Given that NopT negatively regulates rhizobial infection by cleaving LjNFR5 (18) and that NopM dimerization occurs at the plasma membrane (15), we hypothesized that NopM might also associate with NFRs. To test this, we first conducted a luciferase complementation assay in *Nicotiana benthamiana* leaves. A ubiquitin ligase inactive NopM^C338A^ was used in these experiments, because enzymatically active NopM induces cell death in *N. benthamiana* cells (15). The results showed that NopM^C338A^ associated with both LjNFR1 and LjNFR5, as indicated by strong luciferase activity when proteins were co-expressed in *N. benthamiana* leaves (Fig. 2A and 2B). Further validation through a bimolecular fluorescence complementation (BiFC) assay confirmed the interaction between NopM^C338A^ and LjNFR1 or LjNFR5 (Fig. 2C). A co-immunoprecipitation (co-IP) assay provided additional evidence supporting the interaction between NopM^C338A^ and LjNFR1 or LjNFR5 (Fig. 2D, 2E). For a subsequent *in vitro* pull-down assay, the cytosolic domain (CD) of LjNFR1 or LjNFR5 was co-expressed with NopM in *Escherichia coli* cells confirming that NopM physically interactes with both LjNFR1^CD^ and LjNFR5^CD^ (Fig. 2F, 2G).

**Figure 2.**
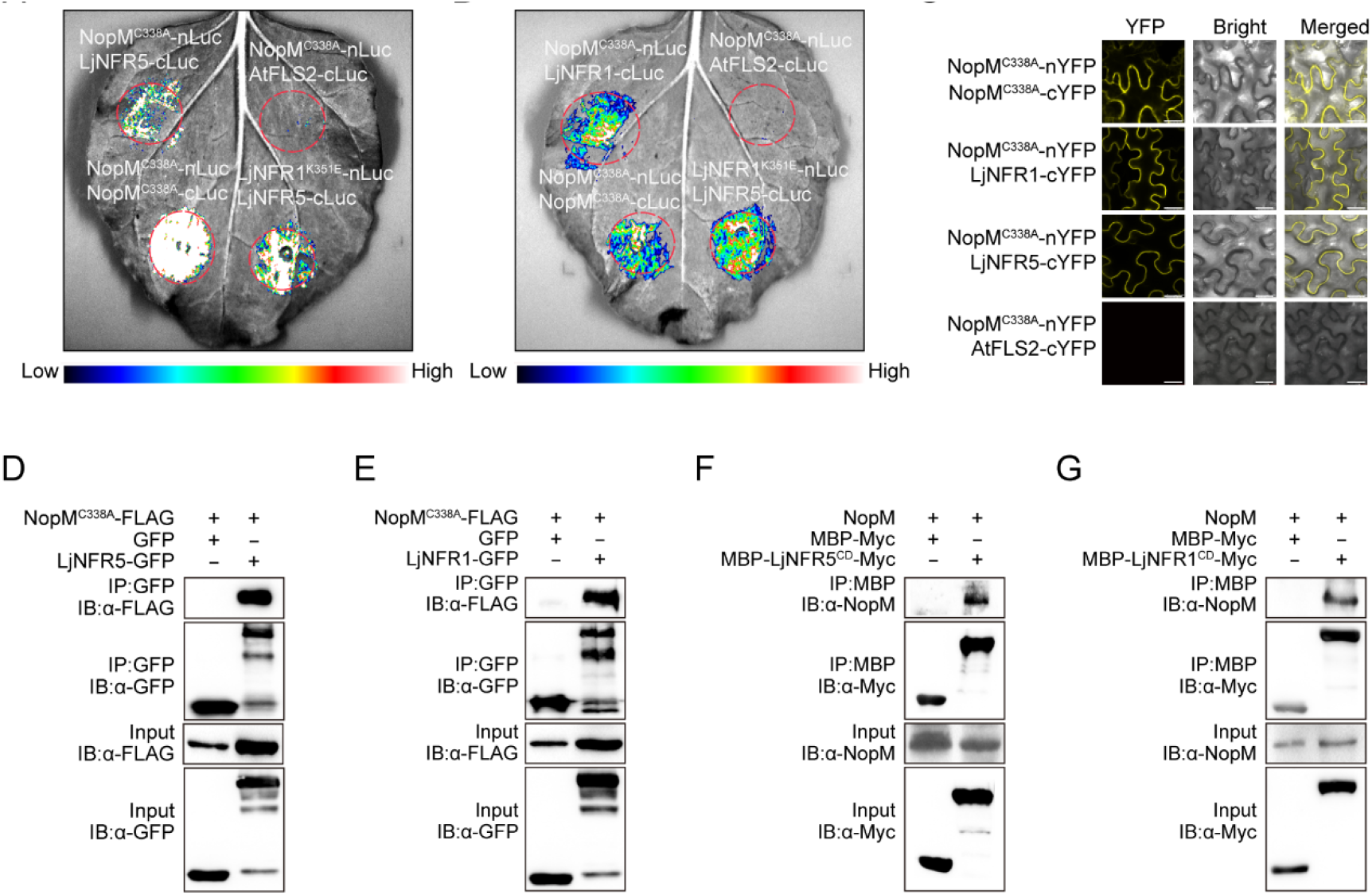
NopM Interacts with LjNFR1 and LjNFR5. (A) and (B) NopM interacts with LjNFR5 or LjNFR1 in split-luciferase assay. NopM^C338A^-nLUC was co-expressed with LjNFR5-cLUC and LjNFR1-cLUC in *N. benthamiana* leaves. NopM^C338A^-nLUC and AtFLS2-cLUC are negative controls, whileLjNFR5-cLUC and LjNFR1^K351E^-nLUC (kinase-dead) served as a positive control. The combination NopM^C338A^-nLUC with NopM^C338A^-cLUC was also analyzed.Luminescence signals indicate protein–protein interactions. (C)NopM interacts with LjNFR5 or LjNFR1 in BiFC assay. NopM^C338A^-nYFP was co-expressed with LjNFR5-cYFP or LjNFR1-cYFP in *N. benthamiana*, and fluorescence was visualized by confocal microscopy. Negative controls included NopM^C338A^-nYFP with AtFLS2-cYFP. The combination NopM^C338A^-nYFP with NopM^C338A^-cYFP was also examined. Bar=25 µm. (D)and (E) NopM interacts with LjNFR5 or LjNFR1 inCo-immunoprecipitation (co-IP) assay. LjNFR5-GFP and LjNFR1-GFP were immunoprecipitated with anti-GFP antibody and detected by immunoblotting with anti-FLAG antibody. (F) and (G) NopM physically interacts LjNFR5 or LjNFR1 in pull-down assay. MBP-LjNFR5-Myc, MBP-LjNFR1-Myc, and NopM were co-expressed in *E. coli* cells. Pull-down was performed with amylose resin and detected by immunoblotting with anti-NopM and anti-Myc antibodies.

### NopM ubiquitinates LjNFR5

Since NopM is an E3 ligase(17), we wondered whether NopM can ubiquitinate NFRs. To test this, a ubiquitination assay was carried out using a reconstituted ubiquitination cascade system (E1, E2, E3 enzymes and ubiquitin) expressed in *E. coli* cells (19). As shown in Fig. 3A, NopM, but not NopM^C338A^, exhibited autoubiquitination activity, which is consistent with previous studies (15). Next, NopM was co-expressed with LjNFR5 in *E. coli* cells. The result showed that NopM could ubiquitinate LjNFR5^CD^ as strong, smeared protein bands were visible on immunoblots. The strong smeared bands of LjNFR5^CD^ suggested that LjNFR5^CD^ might be poly-ubiquitinated (Fig. 3A). Interestingly, NopM did not ubiquitinate LjNFR1^CD^ (Fig. 3B). These findings indicated that NopM selectively ubiquitinates LjNFR5.

**Figure 3.**
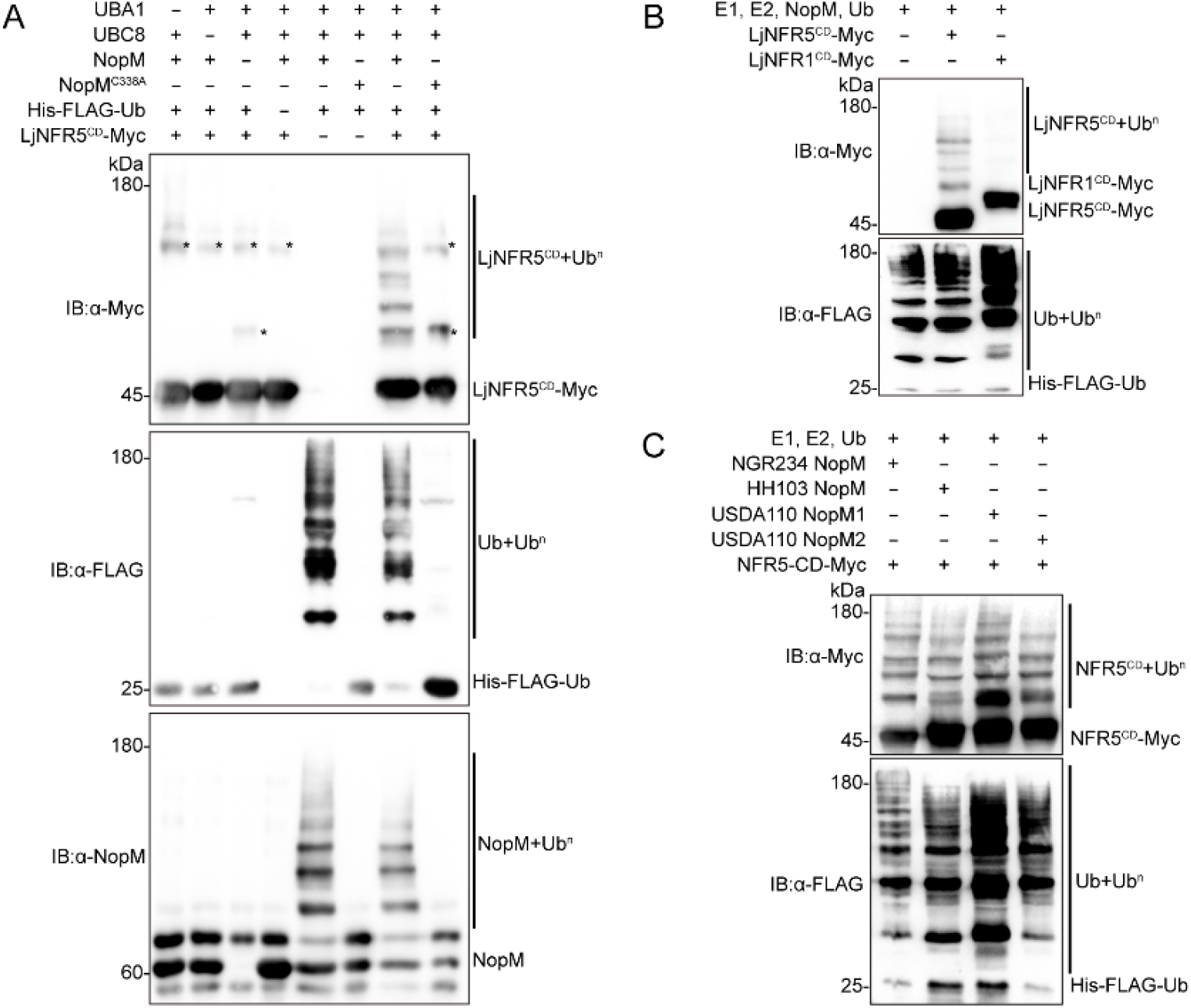
NopM Ubiquitinates LjNFR5. (A)NopM , but not NopM^C338A^, ubiquitinates LjNFR5 in *E. coli*.. NopM or NopM^C338A^, AtUBA1 (E1), AtUBC8 (E2), His-FLAG-Ub, and Myc tagged cytosolic domain (CD) of LjNFR5 (LjNFR5^CD^-Myc) were co-expressed in *E. coli*. Bacterial lysates were subjected to immunoblotting using anti-FLAG, anti-Myc, and anti-NopM antibodies. Asterisks indicated non-specific band. (B)NopM ubiquitinates LjNFR5, but not LjNFR1. LjNFR5^CD^-Myc or LjNFR1^CD^-Myc were co-expressed with NopM, AtUBA1 (E1), AtUBC8 (E2) and His-FLAG-Ub. Western blotting was performed with bacterial lysates using anti-Myc and anti-FLAG antibodies. (C)Different NopM proteins from different rhizobium species ubiquitinate LjNFR5 or GmNFR5. The Myc-tagged cytosolic domain of both NFR5 (NFR5^CD^-Myc), NopM from indicated strains, AtUBA1 (E1), AtUBC8 (E2), and His and FLAG-tagged ubiquitin (His-FLAG-Ub) were co-expressed in *E. coli*. The bacterial lysates were subjected to Western blotting using anti-FLAG and anti-Myc antibodies.

Next, we wondered whether the ubiquitination of LjNFR5 by NopM is a conserved mechanism that also occurs in other symbiotic interactions. We compared NopM sequences from different rhizobium species. Amino acid sequence comparisons showed that the NopM proteins can be divided into three clades (Fig. S4). NopMs of *Bradyrhizobium diazoefficiens* USDA110 and *S. fredii* HH103 were located in Clade I and Clade III, respectively. NopM of *S. fredii* NGR234 was located in Clade II. Two *NopM* genes, named NopM1_USDA110_ and NopM2_USDA110_ were present in the USDA110. *S. fredii* HH103 possesses two NopM genes with identical protein sequences. A multiple sequence alignment indicated that C-terminal E3 ligase domain of NopM was conserved whereas the length of the N-terminal Leucine-Rich Repeats domain was different among the three NopM clades (Fig. S5). We then examined the ubiquitination of soybean GmNFR5^CD^ by NopM1_USDA110_ and NopM2_USDA110_ (Clade I) and NopM_HH103_ (clade III). The results showed that these three NopMs could ubiquitinate GmNFR5^CD^ (Fig. 3C). These findings suggest that NopM homologs from different rhizobium species exhibit a conserved activity in ubiquitinating NFR5 proteins.

### Expression of NopM in *L. japonicus* increases LjNFR5 protein levels

Ubiquitination is an important post-translational modification that controls protein levels by mediating protein degradation or stability (20). To assess whether NopM-mediated ubiquitination regulates LjNFR5 protein levels *in vivo*, we expressed *NopM* and *NopM*^*C338A*^ under the control of a *Ubiquitin* promoter in hairy roots of stable transgenic *L. japonicus* expressing LjNFR5-HA. LjNFR5 was detected at higher levels in NopM-expressing roots than those in roots expressing NopM^C338A^ (Fig. 4A and 4B). These findings indicated that the ubiquitination of LjNFR5 by NopM did not cause degradation, but rather led to increased LjNFR5 protein levels.

**Figure 4.**
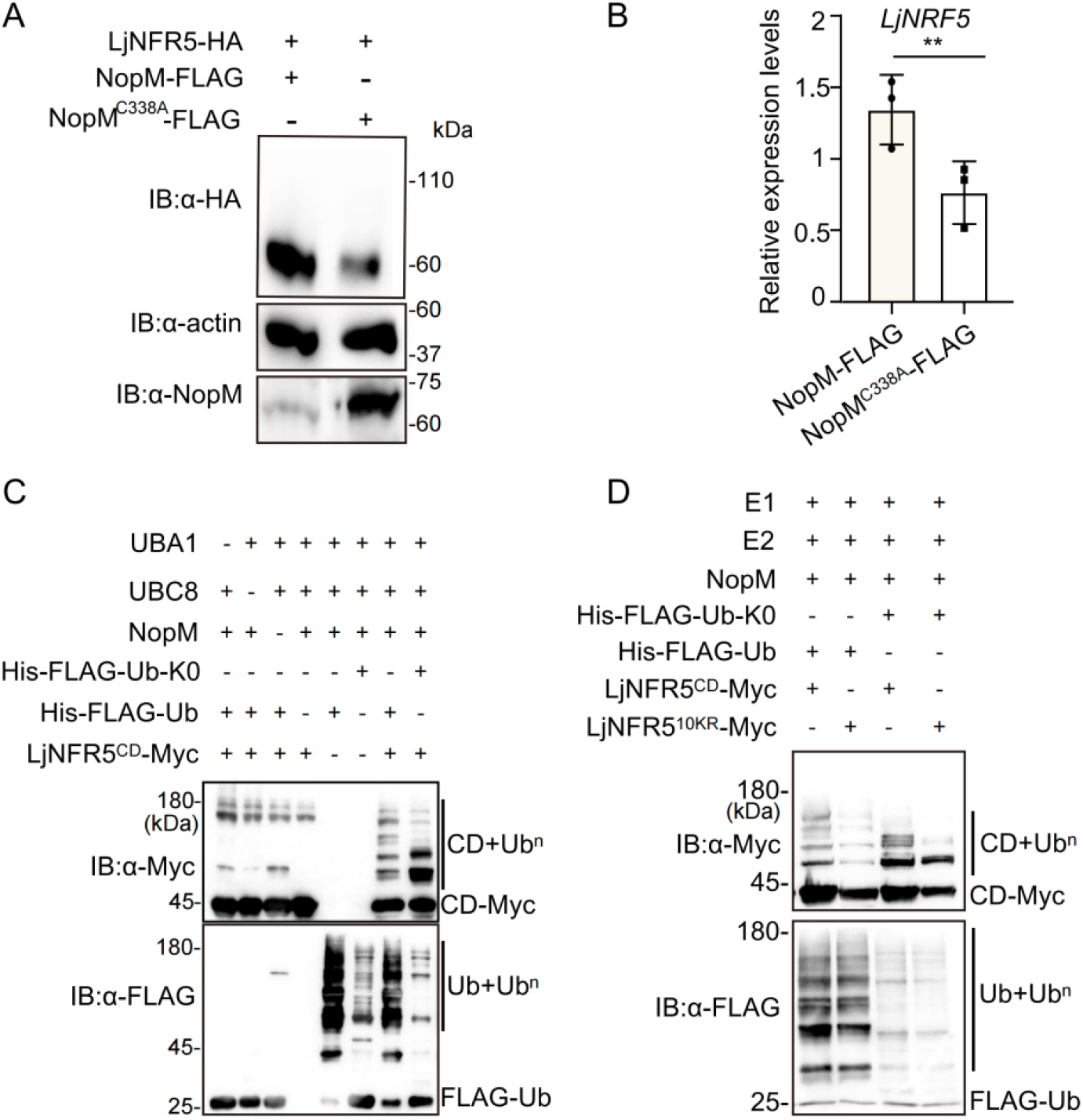
NopM Expression in *L. japonicus* Roots Increases LjNFR5 Levels and NopM-mediated Multi-monoubiquitination of LjNFR5 with Ub^K0^. (A) NopM expression in *L. japonicus* results in increased LjNFR5 protein levels. NopM-FLAG driven by the *Ubiquitin* promoter, was expressed in hairy roots of *proUbiquitin:LjNFR5-HA* plants. Protein levels detected by immunoblotting with anti-HA, anti-NopM, and anti-actin antibodies.(B) Quantification of signals of Western blot bands shown in panel (A). Protein expression levels of LjNFR5 are shown as fold change relative to actin. **p < 0.05 (Student’s t-test).(C) NopM-mediated multi-monoubiquitination of LjNFR5 with Ub^K0^ The ubiquitination assay with *E. coli* cells was performed using His-Flag-tagged Ub^K0^, a ubiquitin variant with all lysine residues substituted to arginine (His-Flag-Ub^K0^) Ubiquitinated proteins were detected by immunoblotting with anti-FLAG and anti-Myc. Asterisks indicated non-specific bands. (D)A similar ubiquitination experiment was performed with LjNFR5^10KR^, in which ten lysine residues are substituted to arginine

Given that ubiquitination of LjNFR5 increased its stability, we proposed that NopM-mediated ubiquitination might form mono-ubiquitination chains. To investigate the type of ubiquitin-chain linkage on LjNFR5, we utilized the Ub^K0^ variant (all lysine residues in ubiqutin were substituted to arginine) (21). Interestingly, the ubiquitination pattern of LjNFR5 in the presence of Ub^K0^ remained similar to that with ubiquitin, suggesting that LjNFR5 might be mono-ubiquitinated by NopM (Fig. 4C). Accordingly, mass spectrometry analysis of LjNFR5 identified ten potential ubiquitinated lysine residues (Fig. S6 and Fig. S7). LjNFR5^10KR^, a mutant version of LjNFR5 with ten lysine residues substituted to arginine residues, still interacts with NopM (Fig. S8A and S8B) and did not abolish the ubiquitination by NopM (Fig. 4D). However, when using the Ub^K0^ variant, LjNFR5 ubiquitination was dramatically reduced (Fig. 4D), suggesting that LjNFR5 still contains additional lysine residues that can be either mono- or poly-ubiquitinated.

To assess the impact of ubiquitinated lysine residues on LjNFR5 function, a cell death test was performed within *N. benthamiana* leaves that depends on co-expression of NFRs. Co-expression of LjNFR1 with LjNFR5 induces rapid cell death as reported previously (22). Co-expression of LjNFR1 with LjNFR5^10KR^ caused a similar cell death response (Fig. S8C). We then generated transgenic hairy roots of *L. japonicus* (Gifu) *nfr5* mutant plants that expressed *LjNFR5* or *LjNFR5*^*10KR*^ driven by the native promoter (*proLjNFR5:LjNFR5, proLjNFR5:LjNFR5*^*10KR*^). When inoculated with GFP-labelled NGR234, hairy roots transformed with the *proLjNFR5: LjNFR5*^*10KR*^ construct showed very few infection events whereas numerous infections were observed in roots expressing LjNFR5 (Fig. S8D). These findings indicate that specific lysine residues required for ubiquitination in LjNFR5 are essential for LjNFR5 to activate symbiotic signaling.

### NopM promotes rhizobial infection and nodule primordia formation

To examine the role of NopM in symbiotic response, we transformed *L. japonicus* MG20 plants with *NopM* driven by *CaMV 35S* or *Ubiquitin* promoters. Unfortunately, we could not obtain stable transgenic plants with these constructs, suggesting that constitutive NopM expression might have a negative effect on plant growth. We therefore generated *L. japonicus* MG20 plants expressing *NopM* driven by the *LjNIN* promoter (*proNIN:NopM*), whose activity is induced during symbiosis (Fig. S9 A-C). Two independent lines were obtained and inoculated with a GFP-tagged *Mesorhizobium loti* MAFF303099 which does not contain *NopT* and *NopM* genes in its genome. As shown in Fig. 5A, 5D and Fig. S9D, increased infection foci and infection threads were observed in these transgenic plants compared to control plants. Consistent with this, transgenic plants also formed more nodule primordia than wild type plants at 14 dpi (Fig. 5B). Finally, to corroborate these results, we generated transgenic roots expressing *NopM* driven by the *Ubiquitin* promoter. Upon inoculation with GFP-tagged *M. loti*, the number of rhizobial infections in these *NopM*-expressing roots was significantly higher than in control roots transformed with the control vector (Fig. 5C). Taken together, these data indicate that NopM positively regulates rhizobial infection and nodule primordia formation in *L. japonicus*.

**Figure 5.**
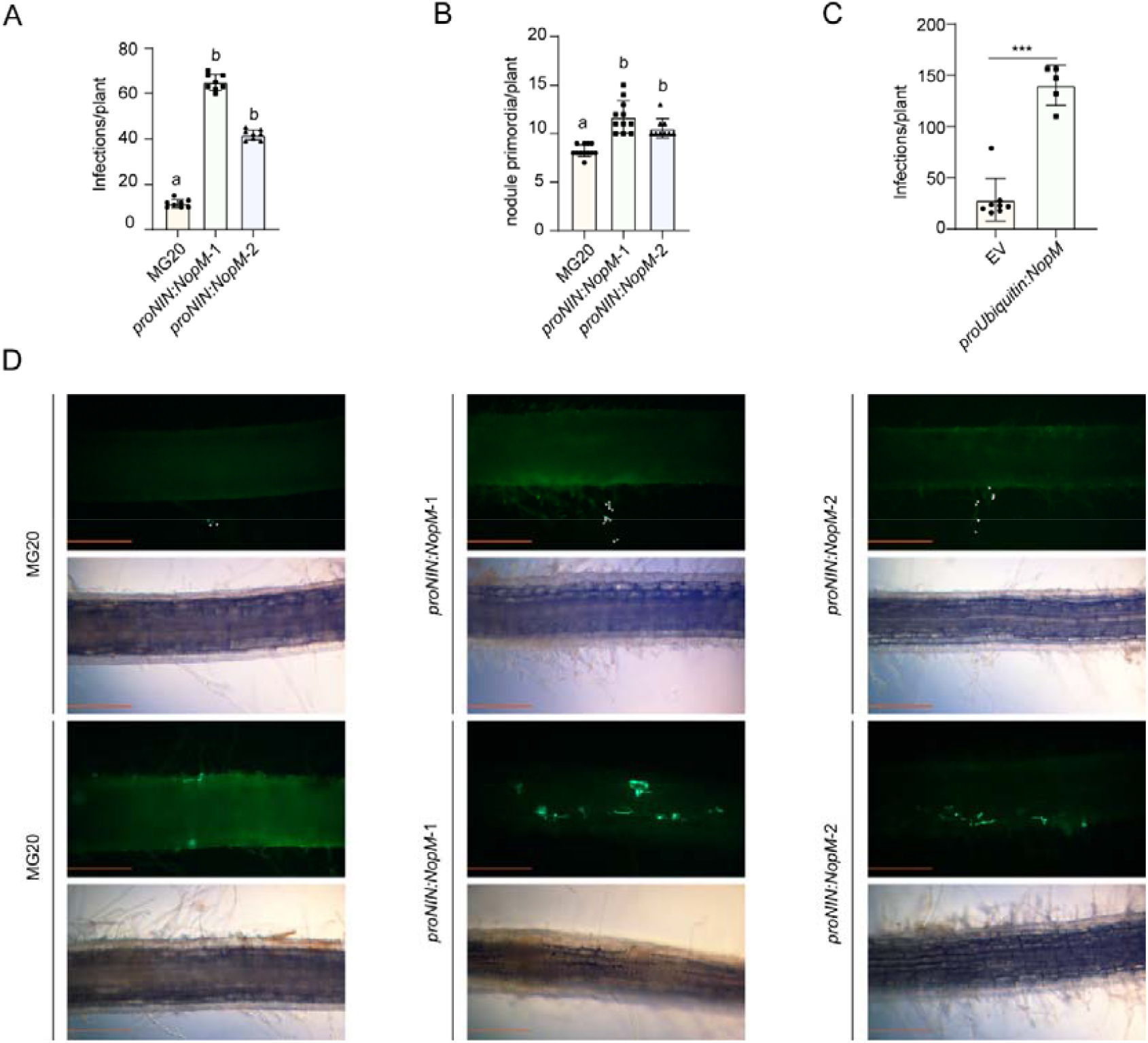
*L. japonicus* Plants Transformed with *proNIN:NopM* and Inoculated with *M. loti* Show Increased Infection and Nodule Primordia Formation. (A) The number of rhizobial infections per plant was significantly greater in transgenic *L. japonicus* MG20 lines carrying the *proNIN:NopM* construct than in the MG20 control plants. Roots were inoculated with GFP-labelled *M. loti* 303099 and microscopically analyzed at 5 dpi. Error bars represent ±SD. ***p < 0.01 (Student’s t-test, n=8). (B) Nodule primordia number per plant were counted in *proNIN:NopM* lines inoculated with GFP-labelled *M. loti* when analyzed at 14 dpi with *M. loti*. Data are mean ±SD (***p < 0.01, Student’s t-test; MG20, n=12; *proNIN: NopM*-1, n=11; *proNIN: NopM*-2, n=9). (C) The number of rhizobial infections per plant increased significantly in hairy roots of *L. japonicus* MG20 carrying the *proUbiquitin:NopM* construct as compared to the empty vector control (EV). Roots were microscopically analyzed at 5 dpi. Error bars represent ±SD. ***p < 0.01 (Student’s t-test; MG20, n=8; *proUbiquitin:NopM*, n=5). (D) Microscopic images illustrating rhizobial root infection events (the upper images were infection foci and the lower images were infection threads) in MG20 wild type and *proNIN:NopM* lines (5 dpi). Fluorescent, GFP-expressing *M. loti* bacteria are shown in the upper images and corresponding brightfield images are presented below. Bar=200 μm. White asterisks indicate bacteria in the upper images.

## Discussion

The study of type III secretion system (T3SS) effectors in gram-negative bacteria has primarily focused on their role in promoting pathogenic infections in plants. While there are parallels between the processes of rhizobial infection and colonization and those of pathogenic bacteria, the specific functions of rhizobial effectors in manipulating the plant symbiotic pathway have remained elusive. This overall aim of study was to unravel the mechanisms by which T3SS effectors regulate rhizobium-legume symbiosis. This research demonstrated that the rhizobial effector NopM can associate with two key receptors, LjNFR1 and LjNFR5, essential for activating symbiotic signaling in plants. NopM selectively ubiquitinates LjNFR5, leading to increased protein levels and enhanced rhizobial infection. These findings suggested a regulatory model in which NopM targets NFR5 in legumes to promote the symbiotic signaling transduction by altering the stability of NFR5.

Root infection by rhizobia often depends on a mixture of T3SS effectors which may act redundantly, synergistically, or antagonistically within host cells. Accordingly, mutation of effector genes show positive, negative or no effects on symbiosis with a given host plant. Analysis of rhizobial double mutants may reveal a symbiotic role of an effector that would not have been visible by mutating a single effector gene. While mutation of *nopT* in NGR234 had a positive effect on infections of *L. japonicus* roots, mutation of other effector genes did not result in an altered phenotype. However, mutation of *nopM, nopP and nopZ* in the *nopT* mutant resulted in reduced infection numbers, indicating that the symbiosis promoting effect of these effector genes was only observed in double mutants. This suggests that NopT and NopM work antagonistically, perhaps targeting the same host protein, to precisely regulate rhizobial infection. The observed role of NopT may be due to interactions with NFRs (18) or protein kinases related to effector-triggered immunity (13, 14). Similar to NopT, we report here that NopM can interact with the LjNFR1 and LjNFR5 of *L. japonicus*. The findings suggest that NopT and NopM play antagonistically redundant roles to fine-tune rhizobial infection in host plants.

LjNFR1 and LjNFR5 play vital roles in perceiving rhizobial NFs, activating symbiotic signaling in *L. japonicus* (2, 22). Once NF signaling is initiated, protein turnover and signaling activity of NFRs may be critical for maintaining proper symbiotic signaling and subsequent rhizobial infection. Previous research demonstrated that mutation in the *LjPUB13* gene suppresses nodulation in *L. japonicus* (23). LjPUB13 is an E3 ligase that interacts with and ubiquitinates LjNFR5 *in vitro* (23). Thus, the biological role of LiPUB13 in promoting rhizobial symbiosis may be similar to that of NopM, as both proteins can ubiquitinate LjNFR5. In general, proteins with K48-linked polyubiquitin chains are rapidly degraded, whereas proteins with other polyubiquitin chains and mono-ubiquitinated proteins are not degraded and often exhibit altered properties (24). In our study, we demonstrate that NopM can mono-ubiquitinate LjNFR5 to stabilize LjNFR5, promoting rhizobial infection in *L. japonicu*s. Hairy roots of *LjNFR5-HA* transgenic plants expressing *proUbiquitin:NopM* showed increased LjNFR5 protein levels when compared to those transformed with *proUbiquitin:NopM*^*C338A*^). Although poly-ubiquitination of LjNFR5 by NopM was detected at weak levels, the mono-ubiquitination of LjNFR5 might be major reason leading to increased protein levels. It is therefore tempting to speculate that NopM preferentially monoubiquitinates LjNFR5 and/or that ubiquitinated LjNFR5 contains polyubiquitin chains without K48-linkages.

Pathogens utilize effectors to facilitate infection by suppressing host immunity, exemplified by the E3 ubiquitin ligase effector AvrPtoB in *Pseudomonas syringae*. AvrPtoB regulates flagellin receptor FLS2 (25) and chitin receptor CERK1 (26) protein stability in *Arabidopsis thaliana*, leading to suppression of PTI. Similarly, NopM may employ a similar strategy to support infection during symbiosis to avoid PTI. NopM dampens the flg22-induced ROS burst in *N. benthamiana* (17) and induces cell death in tobacco *via* its E3 ligase activity (15).

On the other hand, NopM activity in certain plant cells appears to induce responses that are related to ETI: (i) The difficulties we observed in the production of MG20 plants constitutively expressing NopM suggest negative effects of NopM on plant development under certain physiological conditions. (ii) NopM of NGR234 expressed in tobacco induces rapid cell death, which depends on the E3 ubiquitin ligase activity (15). (iii) Mutant analysis showed that NopM of *Bradyrhizobium elkanii* USDA61 triggers ETI-like early nodule senescence in MG20 (27). (iv) Hairy roots of MG20 transformed with a construct containing NGR234 *NopM* of expressed from a tandem *CaMV 35S* promoter formed fewer nodules than plants expressing control vectors or a *NopM*^*C338A*^ construct (15). Thus, although NopM promotes rhizobial infection and nodule primordia formation, the ubiquitin ligase activity of NopM appears to have negative effects on nodulation at later stages of the symbiosis. Host plants may use ETI as a mechanism to avoid unfavorable rhizobia strains.

In NGR234 cultures treated with host flavonoids, genes related to the TTSS are expressed later than *nod* genes required for Nod factor synthesis (28). Thus, it can be assumed that effectors are translocated into host cells in which symbiotic signaling might have already started. Because Nod factor receptors can be viewed as gatekeepers controlling symbiotic signaling transduction, the precise regulation of Nod factor receptors could be the most effective means by which rhizobium fine-tune symbiotic signaling in host cells. NopT of NGR234 was shown to proteolyze LjNFR5 and to inhibit symbiotic infection (18), while in our study NopM positively affected LjNFR5 levels and promoted infection by rhizobia. Mutual regulation on NFR5 by two effectors from the same rhizobium might lead to an optimized or suboptimal infection depending on the host plant.

## Materials and Methods

### Plant Materials and Inoculation with Rhizobia

*Lotus japonicus* Gifu B–129 ecotype was provided by the Center for Carbohydrate Recognition and Signaling (https://lotus.au.dk/) and used for phenotype analysis. *L. japonicus* Gifu *nfr5* mutant (2) and Gifu *proNIN:GUS* transgenic line (29). Wild type plants of *L. japonicus* ‘Miyakojima’ MG20 were used for phenotypic analysis and stable transformation. Transgenic *L. japonicus* MG20 plants overexpressing *LjNFR5* from a *Ubiquitin* promoter (encoding HA-tagged LjNFR5) were constructed in our laboratory and used for hairy root transformation (*proUbiquitin:LjNFR5-HA* line).

All *L. japonicus* seeds were surface-sterilized with H_2_SO_4_ for 6 min, and then with 2 % (w/v) NaClO for 6 min. After synchronization of germination at 4 °C in the dark for 2 days, the seeds were placed on Murashige & Skoog (MS) medium (PhytoTech, M519) containing 1.5 % (w/v) agar for germination (22°C for 2 days in the dark and 2 days in the light). Seedlings were then transferred to growth pots containing vermiculite supplied with ½ B&D (Broughton & Dilworth) medium (30) without nitrogen. Plants were kept under long–day conditions (16 h light/ 8 h dark) at 22 °C. For infection experiments, seven-day-old seedlings were inoculated with *S. fredii* NGR234 and NGR234 mutants or *M. loti* MAFF303099 (OD_600_=0.02).

### Plasmid Construction

All plasmids used in this study were generated using MultiF Seamless Assembly Mix (ABclonal, RK21020). Plasmids pG5XX–HA, pG5XX–Myc, pG5XX–FLAG, pG5XX–nLUC, pG5XX–cLUC, pG5XX–GFP (18), point mutant *LjNFR1*^*K351E*^ and point mutant *NopM*^*C338A*^ using overlapping PCR were generated. To detect the interaction between *LjNFR1, LjNFR5*, and *NopM*^*C338A*^ in *Nicotiana benthamiana* in split-luciferase complementation experiments, the full–length coding sequences of *LjNFR1, LjNFR1*^*K351E*^, *LjNFR5, NopM*^*C338A*^, and *AtFLS2* were cloned into *Xba*I-*Kpn*I digested pG5XX–nLUC (31) and pG5XX–cLUC (31), respectively. For BIFC experiments, the full–length coding sequences of *LjNFR1, LjNFR5, NopM*^*C338A*^, and *AtFLS2* were cloned into pSPYCE (MR) (32) and pSPYNE (R)173 (32), respectively, between *Xba*I and *Bam*HI. For the co-IP assay, the full–length *LjNFR1, LjNFR5, NopM*^*C338A*^ coding sequences were cloned into pG5XX–GFP, pG5XX–FLAG, respectively, between *Xba*I and *Kpn*I. For the pull-down assay with *E. coli*, the sequence encoding the cytosolic domain of *LjNFR5* fused to a Myc tag was cloned into plasmid pMAL-c2X (Addgene, 75286) between *Bam*HI and *Hind*III and the *NopM* coding sequence was cloned into plasmid pACYCDuet (19) between *Sgs*I and *Not*I. The *Myc* tag was cloned from pG5XX–Myc and inserted into plasmid pMAL-c2X between *Bam*HI and *Hind*III, generating plasmid pMAL-c2X-Myc used as control. For *in vitro* ubiquitin assay in *E. coli, NopM* or *NopM*^*C338A*^, and Myc tagged cytosolic domain (CD) of NFR5 (NFR5^CD^-Myc/ LjNFR5^10KR^-Myc) were inserted into plasmid pACYDuet and pCDFDuet (19) between *Sgs*I and *Not*I respectively. His and FLAG tagged ubiquitin (His-FLAG-Ub^K0^) was generated by GenScript Biotech. For hairy roots transformation of *L. japonicus*, the plasmid pUB-RFP (33) was digested with *Kpn*I and *Xba*I, and was inserted fragment *NopM* and *NopM*^*C338A*^. For stable transformation in *L. japonicus NopM* was cloned into *Kpn*I-*Xba*I digested pUB-Hyg-FLAG (33), generating plasmid pUB-Hyg-NopM-FLAG with a *Ubiquitin* promoter. A 3-kb promoter fragment upstream of the start codon of *NIN* was then cloned from genomic *L. japonicus* MG20 DNA and inserted into *Xba*I-*Pst*I digested pUB-Hyg-NopM-FLAG to replace the *Ubiquitin* promoter. The resulting plasmid was named proNIN-NopM-Hyg-FLAG. All primers used are listed in the Supplementary Table S1.

### Stable Transformation in *L. japonicus*

The plasmid proNIN-NopM-Hyg-FLAG was transformed into *L. japonicus* MG20 seedlings using *Agrobacterium tumefaciens* strain EHA105 as described by Handberg and Stougaard (34). 50 mg/ml hygromycin B (Sangon Biotech, A600230) was used for selection of transgenic plants. The final positive transgenic plants were detected by PCR analysis.

### Transient Expressing in *N. benthamiana* Leaves

*A. tumefaciens* EHA105 strains carrying various constructs were cultivated overnight and resuspended with an *Agrobacterium* culture harboring the P19 suppressor with the ratio 1:1 in infiltration buffer (10 mM MgCl_2_, 10 mM MES pH=5.8 and 100 μM acetosyringone). The bacterial suspensions were infiltrated into leaves of 4–week–old *N. benthamiana* plants. Leaves were harvested for two days post infiltration.

### BiFC and Split Luciferase Assays

In the BiFC assay with transformed *N. benthamiana* leaves, the eYFP fluorescence of fusion proteins was analyzed 2–3 days after infiltration using a confocal microscope (Leica TCS SP8).

In the split–luciferase assay, the plasmids were transformed into *N. benthamiana* leaves using *Agrobacterium* strain EHA105. Three days after infection, leaves were sprayed with 1□mM luciferin (Promega, E1603) and 0.02 % (v/v) Triton X–100. The LUC activity was detected using the Tanon Imaging system (Tanon, 4600).

### Co-IP and Pull-down Assays

For the co-IP assay, total proteins were extracted from the *N. benthamiana* leaves in IP buffer (50 mM Tris-HCl, pH=7.4, 150 mM NaCl, 1□mM EDTA, 1 % TritonX-100, 5 % glycerol) supplemented with a protease inhibitor cocktail (Yuanye, S25910). Samples were incubated at 4 °C for 10 min and centrifuged at 12,000 × g for 10□min at 4 °C. The supernatant was added to 10□μL anti-GFP Nanobody Agarose Beads (Alpa Life Bio, KTSM1301) and incubated at 4 °C for 3□h. The beads were washed at least five times with washing buffer (50 mM Tris-HCl, pH=7.4, 150 mM NaCl, 1 mM EDTA). Western blotting was detected by anti-GFP (ABclonal, AB_2770402) and anti-FLAG (Sigma, F1804) antibodies.

For the pull-down assay with *E. coli* BL21 (DE3), protein synthesis of co-expressed MBP-LjNFR5^CD^-Myc or MBP-Myc and His-NopM was induced with 0.5 mM IPTG and incubation at 28 °C for 18 h. MBP-LjNFR1^CD^-Myc, or MBP-Myc and His-NopM were co–expressed in a similar way but bacteria were kept at 18 °C for 20 h. Total proteins were extracted from *E. coli* in column buffer (20 mM Tris-HCl, pH=7.4, 0.2□M NaCl, 1 mM EDTA). Samples were centrifuged at 12,000 × g for 10□min at 4 °C. The supernatant was added to 20□μL amylose resin beads (New England Biolabs, E8022S) and incubated at 4 °C for 3□h. The beads then were washed at least five times with column buffer. Western blotting was detected by anti-Myc (Bio–Legend, 626808) antibody and anti-NopM antibody kindly provided by Dr. Xin from North East Agriculture University.

### *In Vitro* Ubiquitination Assay

The *in vitro* ubiquitination assay was performed using genes encoding AtUBA1 (E1), AtUBC8 (E2), NopM (or NopM^C338A^), ubiquitin (or His-FLAG-Ub^K0^) and the cytosolic domain of LjNFR5 (or LjNFR1) as described previously (19). Different combinations of vectors were transformed into the *E. coli* BL21 (DE3). Recombinant protein expression was induced with 0.5 mM IPTG and cells were grown at 28 °C for 20 h. The cell lysates were subjected to Western blotting analysis using anti-Myc (Bio–Legend, 626808) antibody, anti-FLAG (Sigma, F1804) antibody, and anti-NopM antibody.

### Generation of Hairy Roots Expressing NopM and NopM^C338A^ in *L. japonicus*

The pUB-RFP vectors carrying *NopM* and *NopM*^*C338A*^ were transformed into *proUbiquitin:LjNFR5-HA* seedlings and Gifu seedlings using *A. rhizogenes* LBA1334 (33). The *proLjNFR5:LjNFR5-GFP* and *proLjNFR5:LjNFR5*^*10KR*^*-GFP* vectors were transformed into Gifu *nfr5* seedlings using *A. rhizogenes* LBA1334. The expression of RFP and GFP markers enabled selection of transgenic hairy roots using a fluorescence stereo microscope (Nikon, SMZ18).

### Analysis of proteins expressed in *L. japonicus*

Total proteins were extracted from transgenic hairy roots using IP buffer (20 mM Tris–HCl pH=7.5, 50 mM NaCl, 1 mM EDTA, 0.1 % SDS, 1 % Triton X-100) supplemented with a protease inhibitor cocktail (Yuanye, S25910). Western blotting was performed with anti-HA–peroxidase (Sigma, clone 3F10), anti-actin (ABclonal, AC009), and anti-NopM antibodies.

## Acknowledgments

We thank Jens Stougaard (Aarhus University) for providing *nfr5* mutant and *proNIN:GUS* transgenic seeds. We also thank Dongping Lv (Shanghai Jiaotong University) for providing bacterial ubiquitination system strains. We also thank Dr. Gar Stacey for his valuable comments on this manuscript. This work was supported by the National Key R&D Program of China (2019YFA0904700), the National Natural Science Foundation of China (32090063), and a Self-Innovation grant from the National Laboratory (AML2023B01). The qPCR and microscopy data were acquired from the Core Facility Center run by the National Key Lab of Agricultural Microbiology. Mass spectrometry analysis was performed by the Center for Protein Research at Huazhong Agricultural University.

## Notes

### Competing Interest Statement

The authors have declared no competing interest.

### Summary of Updates

Dear Editors, In the last month, we have made some revisions to refine the manuscript and eliminate any typographical errors. These changes have significantly enhanced the quality of the manuscript, and thus, we would like to submit the updated version. Many thanks, Yangrong

