## Supplemental Figures for "Rhizobial effector NopM ubiquitinates Nod factor receptor NFR5 and promotes rhizobial infection in *Lotus japonicus*"

The following Supporting Information is available for this article:

**Fig. S1** **Schematic of NGR234 Mutants.**

Schematic representation of constructed *S. fredii* NGR234 mutant derivatives and double mutants with a ΩKan interposon (1.6-kb *proKan:Kanamycin* cassette) inserted at nucleotide position 18-25 bp of a given gene.

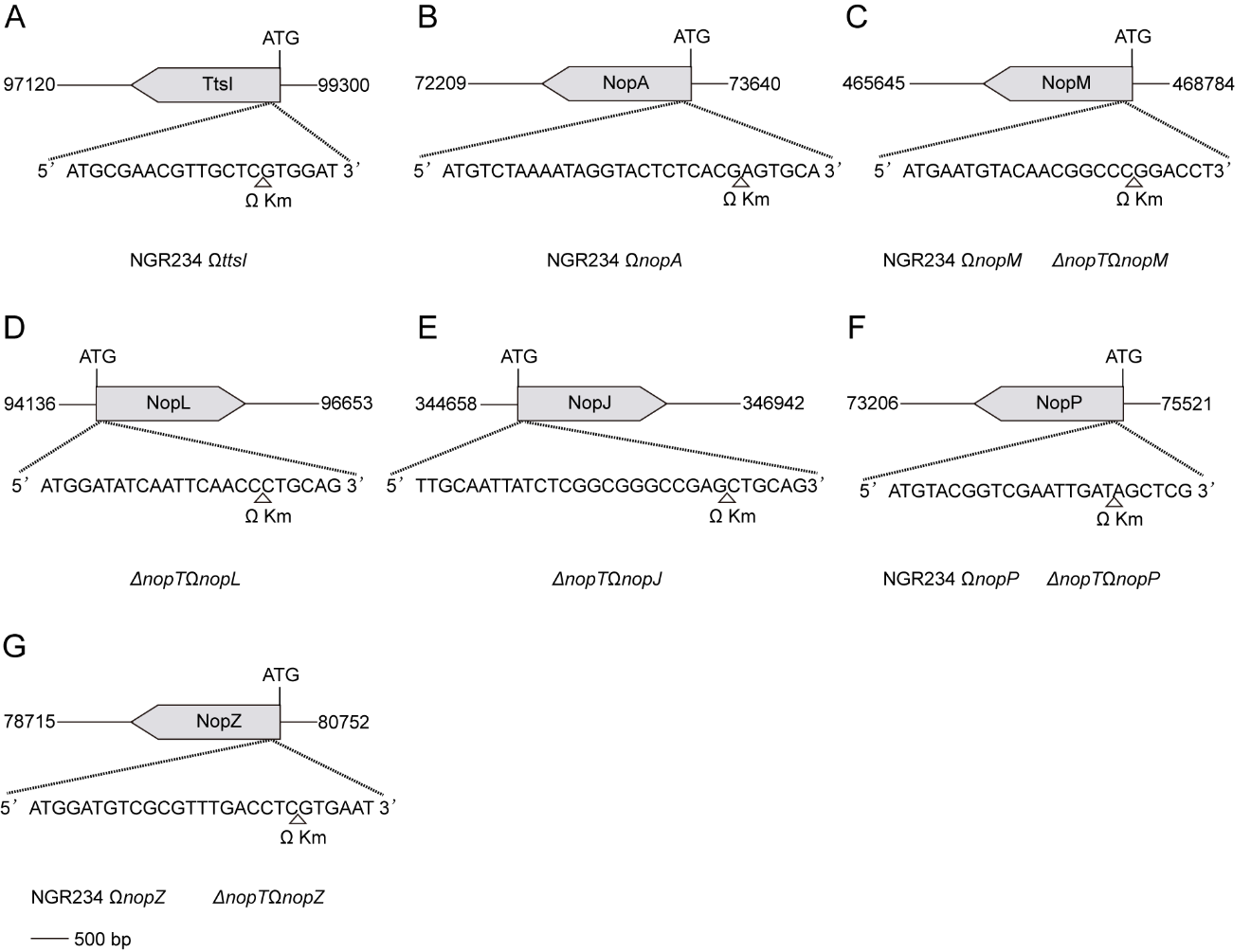

**Fig. S2** **PCR Analysis of NGR234 Mutants.**

PCR confirmation of NGR234 mutants used in this study. Mutants show an increased amplicon length (amplified full-length sequence; “full”) due to insertion of an interposon (1.6-Kb ΩKan).

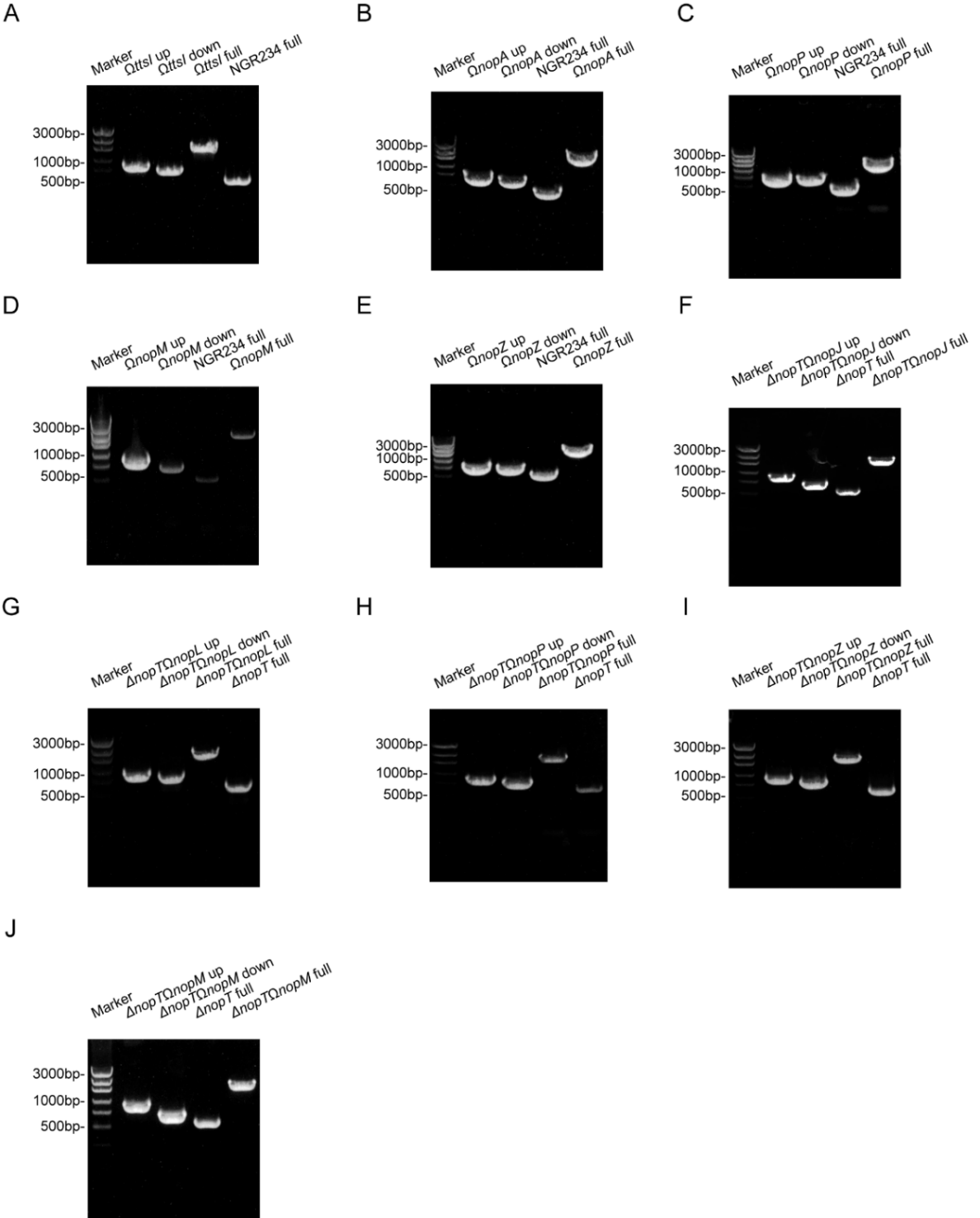

**Fig. S3** **GUS Staining Pictures for Roots of *L. japonicus* Inoculated with *S. fredii* NGR234 and Mutant Strains.**

Roots of *L. japonicus* Gifu expressing *GUS* under the control of *pNIN* (Gifu *pNIN:GUS*) were inoculated with NGR234 and indicated mutants. Roots were stained with X-Gluc solution at 1 dpi. Infection sites showing GUS activity are shown in blue. Pictures in different panels indicate different experiments. Bar=500 μm.

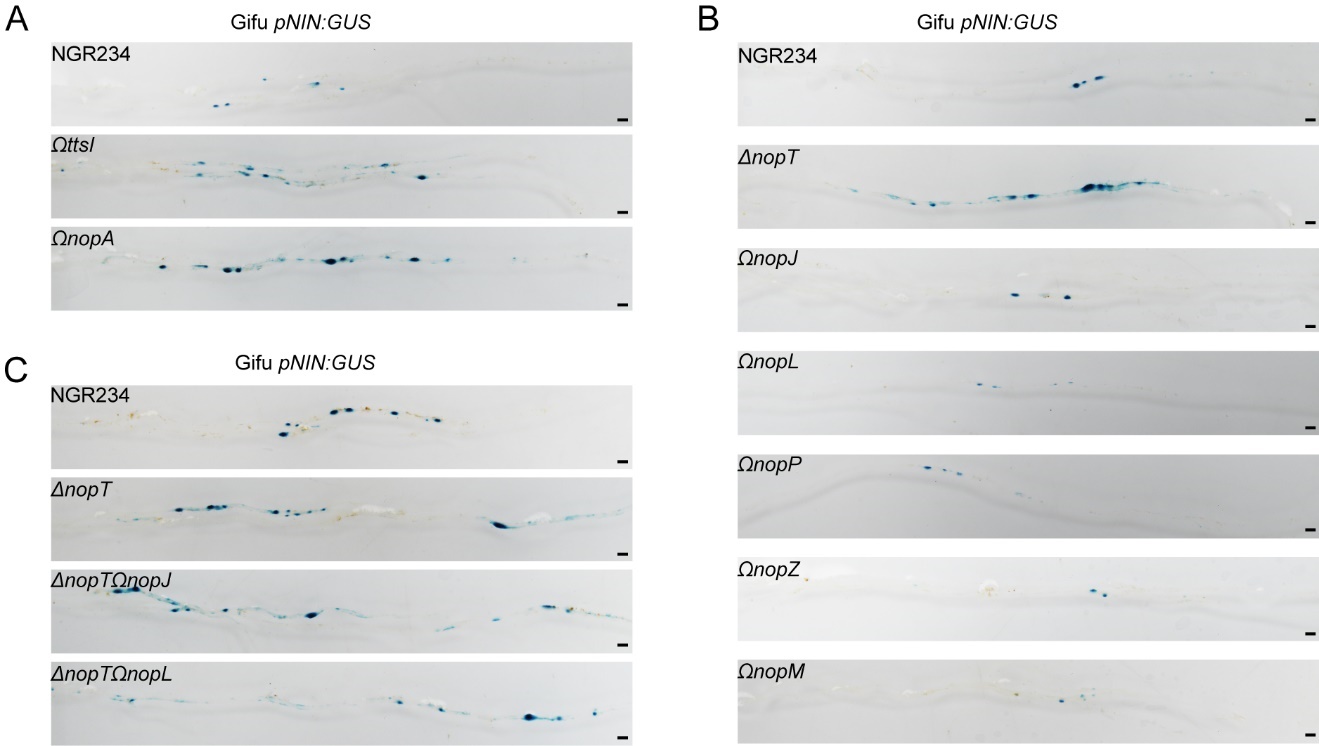

**Fig. S4** **Phylogenetic Tree of NopM Homologs in Different Rhizobial Species.**

A phylogenetic tree based on the amino acid sequence of NopM homologs from different rhizobial species was constructed using the neighbor-joining method. Bootstrap values based on 1000 replications are listed as percentages at branching points. The bar indicates a distance of 0.2 substitutions per amino acid position in the alignment. The homologs were grouped into three clades.

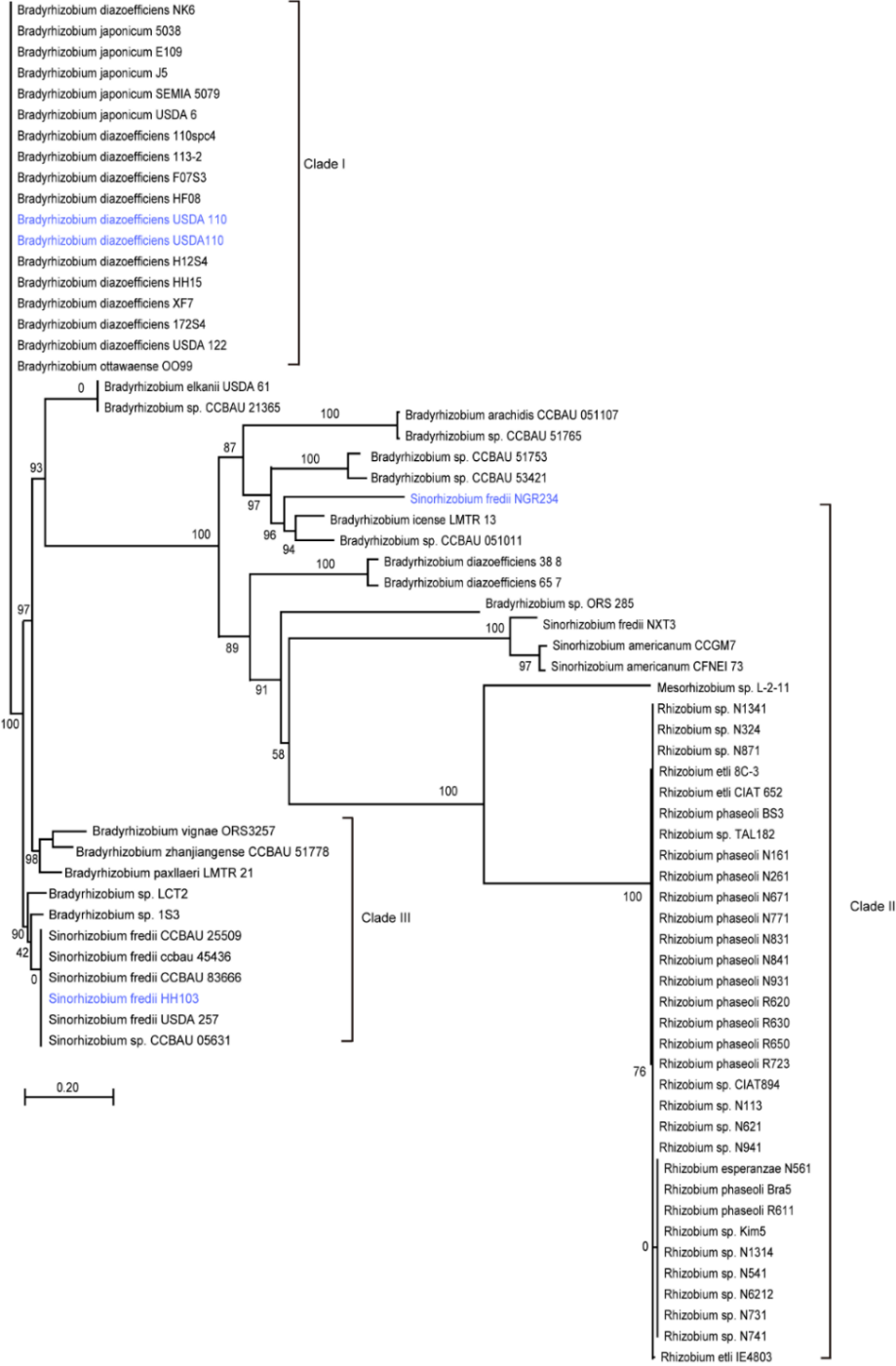

**Fig. S5 Amino Acid Sequence Alignment of NopM Homologous in Different Rhizobial Species.**

The multiple sequence alignment tool MultAlin was used to align NopM protein sequences from different rhizobial species. Conserved amino acids are labelled in red.

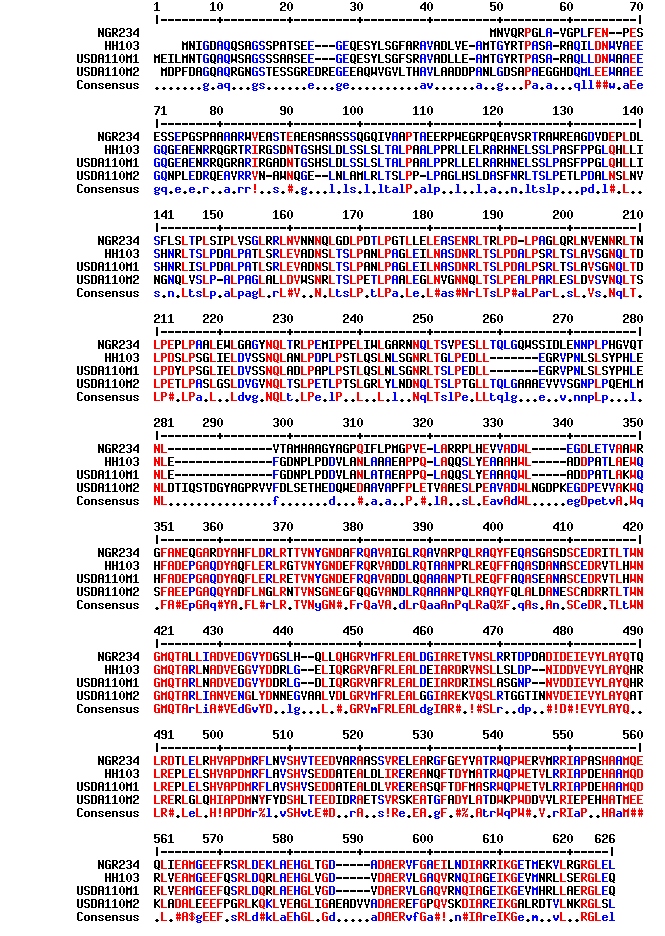

**Fig. S6** **Amino Acid Sequence of LjNFR5 with Identified Lysine Ubiquitination Sites.**

The full-length amino acid sequence of LjNFR5 is depicted. The cytosolic domain (CD) of LjNFR5 is labeled in blue and the identified lysine ubiquitination sites are marked in red.

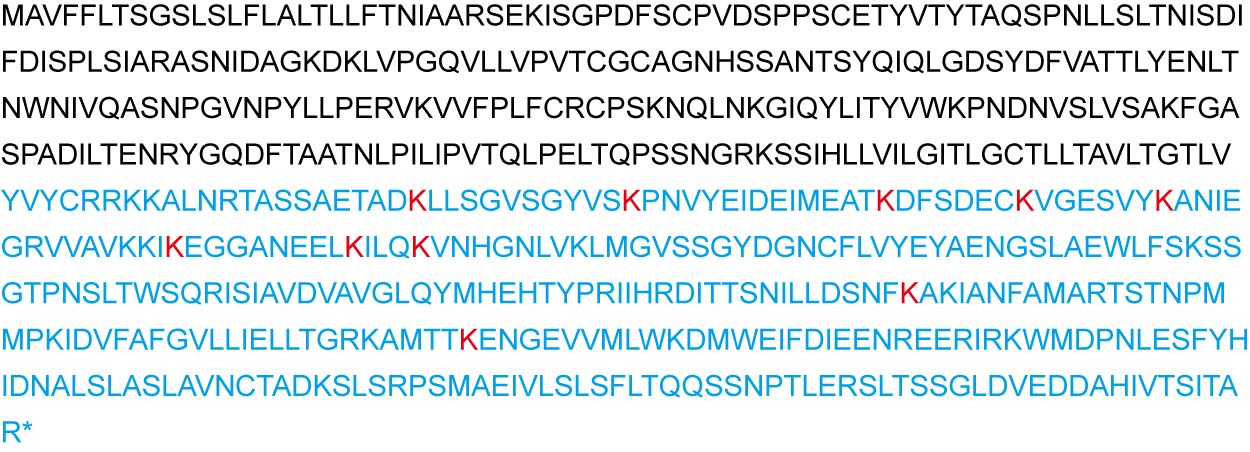

**Fig. S7** ***In Vitro* Ubiquitination Sites of LjNFR5 Identified by Mass Spectrometry.**

Ubiquitination sites in LjNFR5 identified by mass spectrometry. MS/MS spectra of peptides containing ubiquitinated lysine residues in LjNFR5 are shown in (A) K289, (B) K300, (C) K314, (D) K321, (E) K328, (F) K342, (G) K351, (H) K355, (I) K446, and (J) K488. The mass spectrometry analysis was performed once.

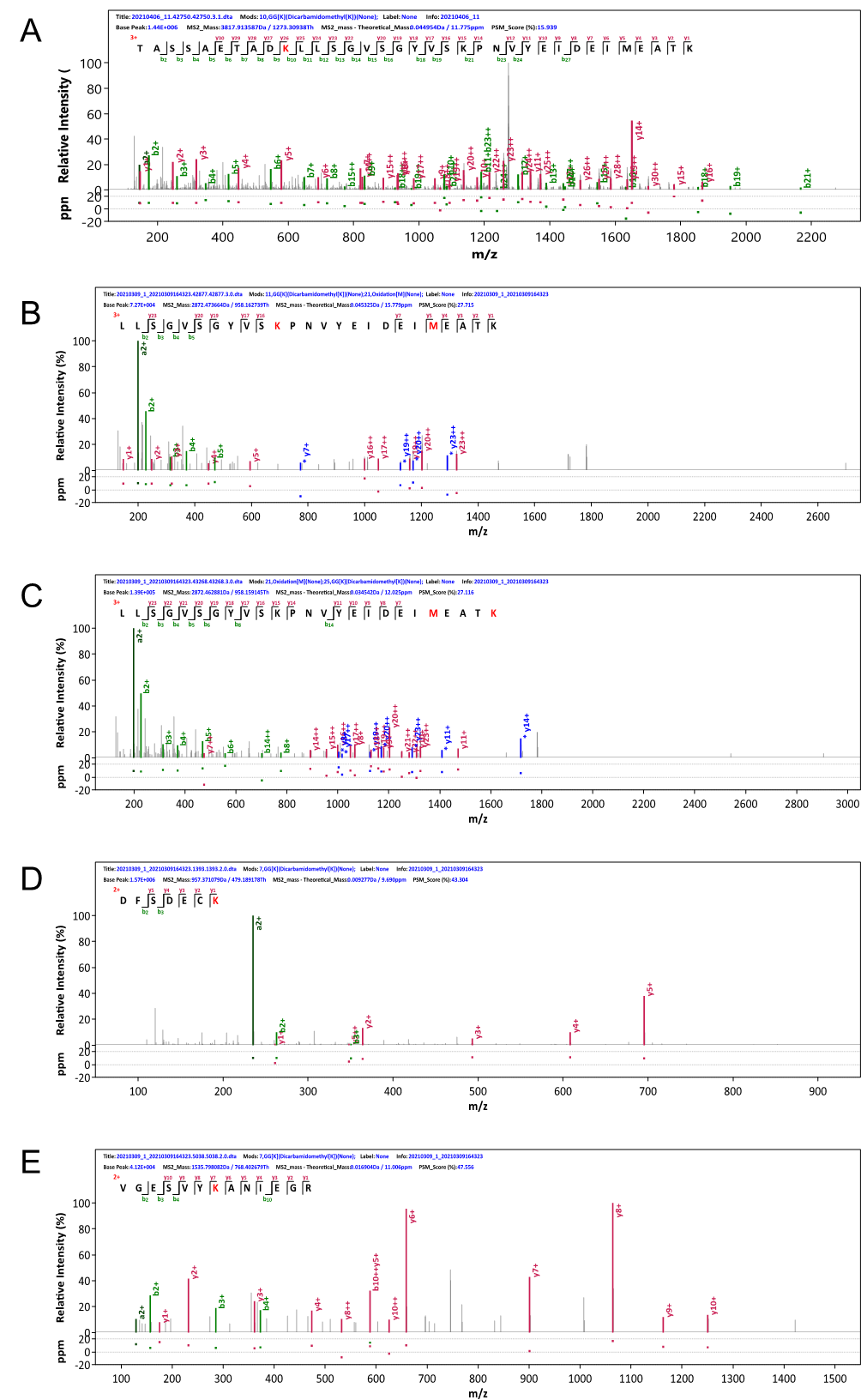

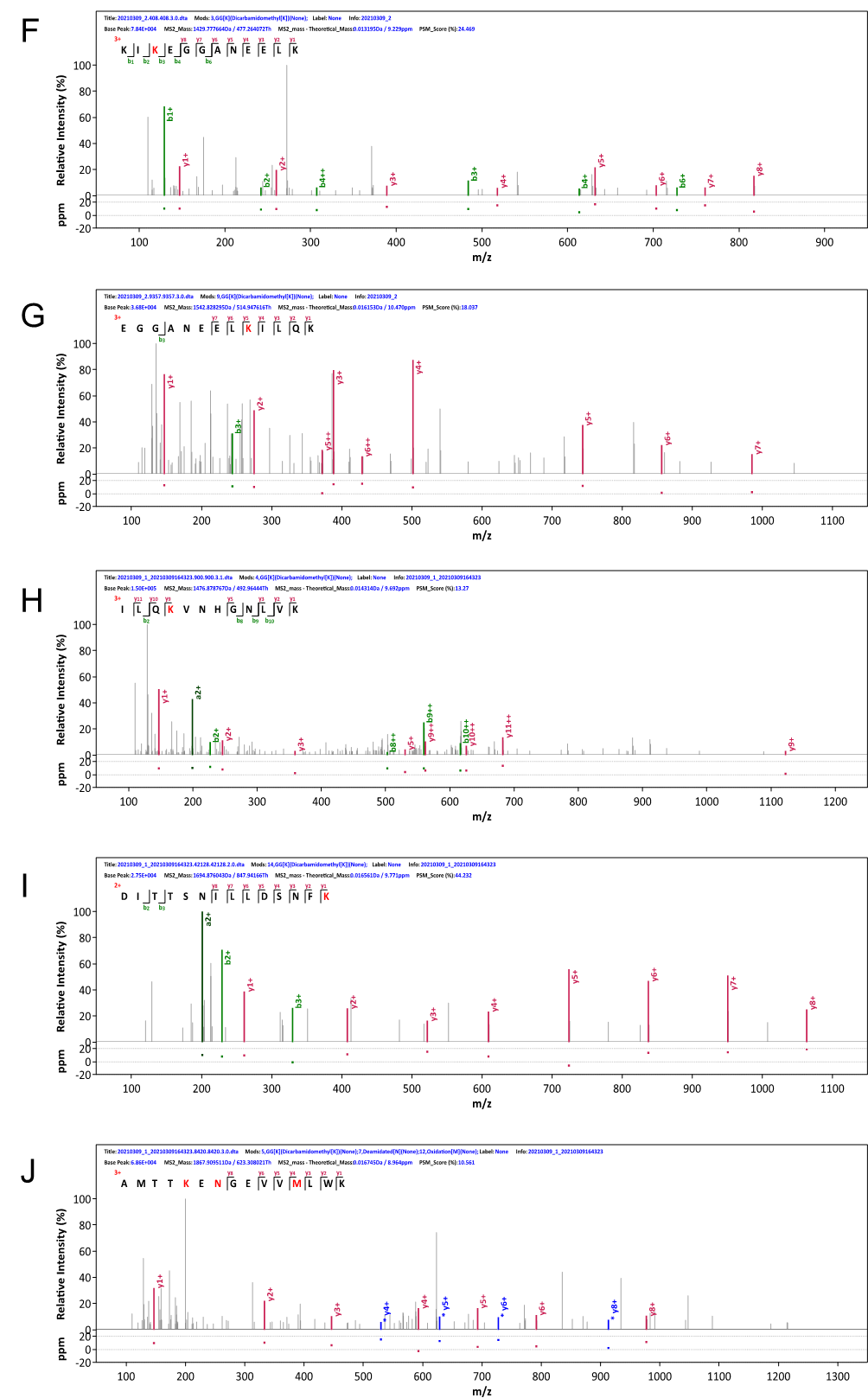

**Fig. S8** **LjNFR5^10KR^ Does Not Affect Interactions with Nod Factor Receptors but Is Symbiotically Inactive.**

(A) NopM interacts with LjNFR5^10KR^ in the split-luciferase complementation test. NopM-nLUC and LjNFR5^10KR^-cLUC were co-expressed *in Nicotiana benthamiana*. Luminescence signals represent protein–protein interactions.

(B) NopM directly interacts with LjNFR5^10KR^ in the pull-down assay. MBP-LjNFR5^10KR^-Myc and NopM were co-expressed in *E. coli* cells. Proteins were subjected to a MBP based pull-down assay and NopM protein was detected by immunoblotting with the anti-NopM antibody.

(C) LjNFR5^10KR^ and LjNFR1 expressed in *N. benthamiana* induce cell death. LjNFR1 and LjNFR5 were used as positive controls. Leaf discs expressing indicated proteins are marked by red circles. The picture is representative of at least three independent biological replicates.

(D) Number of infections per plant in hairy roots of the *L. japonicus* Gifu *nfr5* mutant expressing *LjNFR5* or *LjNFR5^10KR^* driven by the native promoter (*proLjNFR5:LjNFR5* and *proLjNFR5: LjNFR5^10KR^*). Plants were inoculated with *S. fredii* NGR234 and harvested at 7 dpi. Values indicate mean ± SD (Student’s t test: ***p < 0.01; significant difference compared with the respective control).

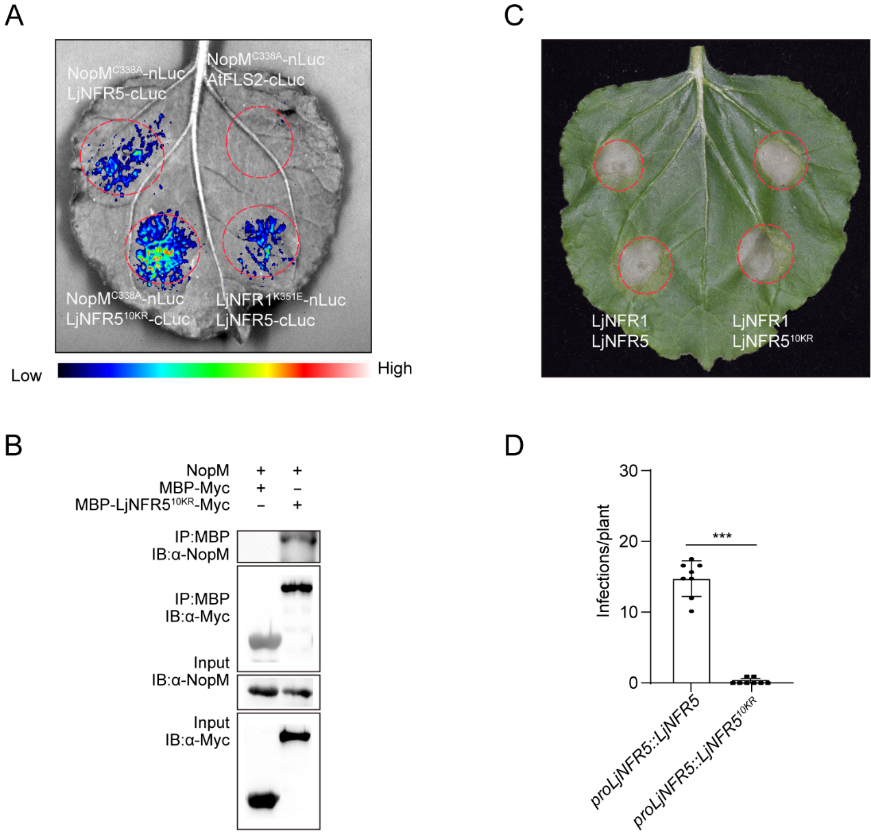

**Fig. S9** **Detection of *NopM* DNA, *NopM* Expression and NopM Protein in Transgenic Plants.**

(A) PCR analysis of *NopM* in *L. japonicus* MG20 and in transgenic *proNIN:NopM* plants. *LjNFR5* was used as the reference gene.

(B) NopM protein levels in these plants were detected by immunoblotting with the anti-NopM antibody. Plants were inoculated with *M. loti* MAFF303099 and harvested at 48 h dpi.

(C) Expression analysis of *NopM* and *ATPase* genes in *proNIN:NopM* plants by quantitative RT-PCR. *L. japonicus* MG20 served as a negative control. The plants were inoculated with *M. loti* MAFF303099 and harvested at 48 h dpi. Values indicate mean ± SD (n=3).

(D) Rhizobial infection in MG20 and *proNIN:NopM* plants inoculated with GFP-labelled *M. loti* MAFF303099. Plants were harvested at 5 dpi. Pictures were enlarged from Fig.4D. Fluorescent, GFP-expressing bacteria are shown in green. Bar=50 μm. Curled root hairs are marked by asterisks.

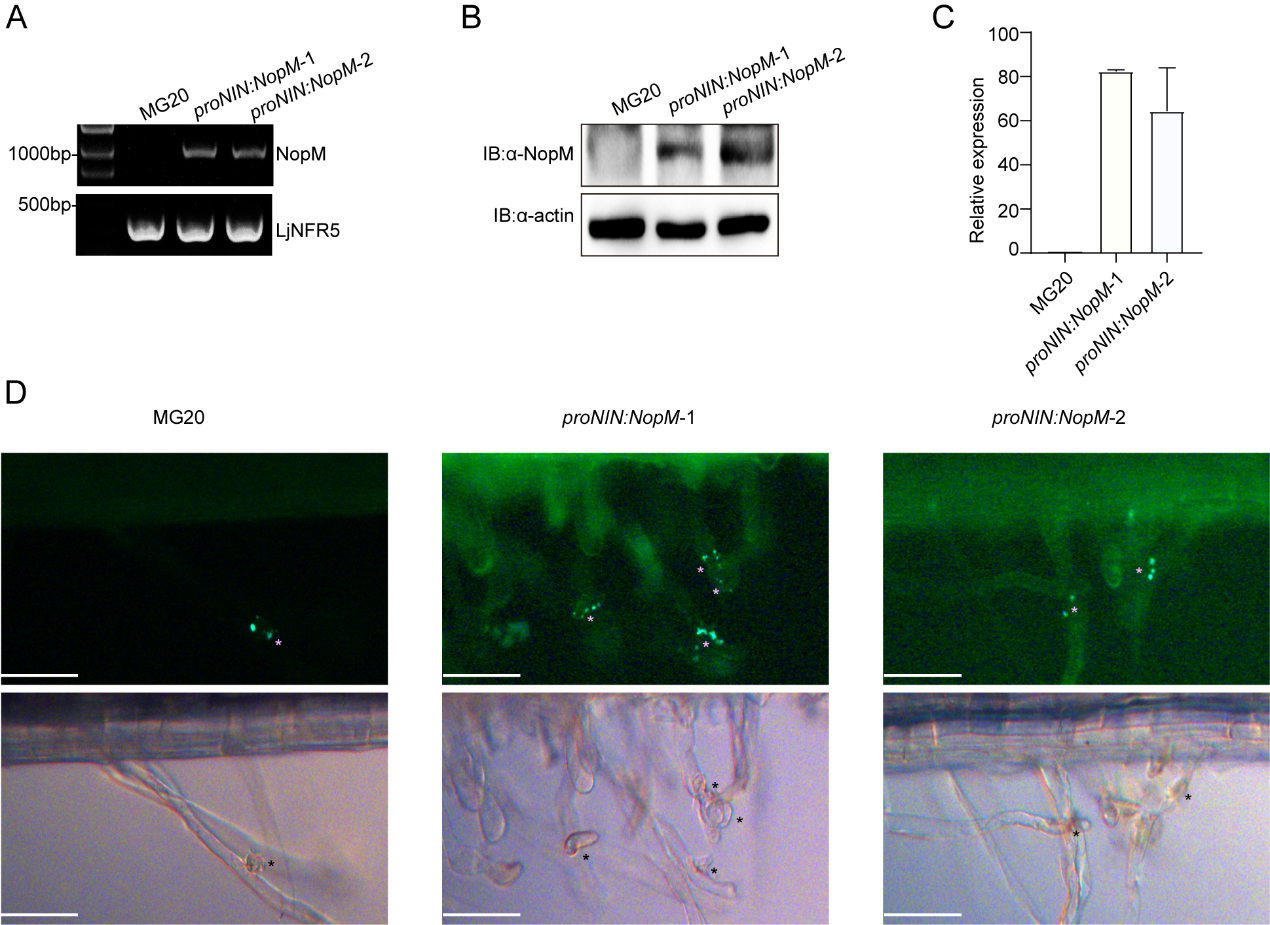
