## Supplemental Table for "Rhizobial effector NopM ubiquitinates Nod factor receptor NFR5 and promotes rhizobial infection in *Lotus japonicus*"

The following Supporting Information is available for this article:

**Table S1** **Primers Used in This Study.**

| Name | Sequence (from 5’ to 3’) | Usage |
| --- | --- | --- |
| pJQ200SK-Kan-F | CTTGATATCGAATTCCTGCAGATGAGCCATATTCAACGGG | Cloning of ΩKan |
| pJQ200SK-Kan-R | CCGCTCTAGAAcTAGTGGATCCTTAGAAAAACTCATCGAGCATCAA | Cloning of ΩKan |
| pJQ200SK-Paph-F | CTTGATATCGAATTCCTGCAGAGCTCGCACGCTGCC | Cloning of the Paph promoter |
| pJQ200SK-Paph-R | GAATATGGCTCATCTGCAAATTATATCTCCTTCTTATTTCTAGTCAAGA | Cloning of the Paph promoter |
| pJQ-Kan-ttsI-up-F | CTTGATATCGAATTCCTGCACCGAAGAATCGATTTGCTC | Homologous fragment |
| pJQ-Kan-ttsI-up-R | CGTGCGAGCTCTGCACTAGAGCAACGTTCGCATTTCA | Homologous fragment |
| pJQ-Kan-ttsI-down-F | GATGAGTTTTTCTAAGGATCCACTAgTGTGGATACGGATCTTACCCG | Homologous fragment |
| pJQ-Kan-ttsI-down-R | GCGGCCGCTCTAGAAcTAGTTGATGACGCGTCAGTCAATT | Homologous fragment |
| pJQ-Kan-nopA-up-F | GTACCGGGCCCCCCCTCGAGcacgtttatatgcaatagcgct | Homologous fragment |
| pJQ-Kan-nopA-up-R | CGTGCGAGCTCTGCAGcgtgagagtacctattttagacatgt | Homologous fragment |
| pJQ-Kan-nopA-down-F | TTGATGCTCGATGAGTTTTTCTAAGGATCCagtgcagttggagctgg | Homologous fragment |
| pJQ-Kan-nopA-down-R | GCGGTGGCGGCCGCTCTAGAgtctcatcatcagtgcgattat | Homologous fragment |
| pJQ-Kan-nopJ-up-F | GTACCGGGCCCCCCCTCGAGagccagttcagccgtca | Homologous fragment |
| pJQ-Kan-nopJ-up-R | GCAGCGTGCGAGCTCTGCAGctcggcccgccgA | Homologous fragment |
| pJQ-Kan-nopJ-down-F | ATGCTCGATGAGTTTTTCTAAGGATCCattggaaatccgtcgagcc | Homologous fragment |
| pJQ-Kan-nopJ-down-R | GCGGTGGCGGCCGCTCTAGAcgcgcctgctggt | Homologous fragment |
| pJQ-Kan-nopL-up-F | GTACCGGGCCCCCCCTCGAGctcctgtcgctcgtctc | Homologous fragment |
| pJQ-Kan-nopL-up-R | CGTGCGAGCTCTGCAGggttgaattgatatccatcctgtttctcc | Homologous fragment |
| pJQ-Kan-nopL-down-F | TTGATGCTCGATGAGTTTTTCTAAGGATCCagcccactaaacgc | Homologous fragment |
| pJQ-Kan-nopL-down-R | GCGGTGGCGGCCGCTCTAGAgcacgaacgccgctatc | Homologous fragment |
| pJQ-Kan-nopP-up-F | GTACCGGGCCCCCCCTCGAGatatccgtgcgtgcaga | Homologous fragment |
| pJQ-Kan-nopP-up-R | CGTGCGAGCTCTGCAGatcaattcgaccgtacatggtga | Homologous fragment |
| pJQ-Kan-nopP-down-F | TTGATGCTCGATGAGTTTTTCTAAGGATCCagctcgtccgatttccat | Homologous fragment |
| pJQ-Kan-nopP-down-R | GCGGTGGCGGCCGCTCTAGAgtcacatgaagtcatcttcgtaagtct | Homologous fragment |
| pJQ-Kan-nopZ-up-F | GTACCGGGCCCCCCCTCGAGgcgagataggtcttgctgC | Homologous fragment |
| pJQ-Kan-nopZ-up-R | GCGGCAGCGTGCGAGCTCTGCAGgaggtcaaacgcga | Homologous fragment |
| pJQ-Kan-nopZ-down-F | TGCTCGATGAGTTTTTCTAAGGATCCgtgaattatgcttacgcttca | Homologous fragment |
| pJQ-Kan-nopZ-down-R | GCGGTGGCGGCCGCTCTAGAcggccgagagagcga | Homologous fragment |
| pJQ-Kan-nopM-up-F | CTTGATATCGAATTCCTGCAGCTTCACATGTCCGAG | Homologous fragment |
| pJQ-Kan-nopM-up-R | CGTGCGAGCTCTGCAGGGGCCGTTGTACATTCATCG | Homologous fragment |
| pJQ-Kan-nopM-down-F | GATGAGTTTTTCTAAGGATCCACTAgTCCCGGACTTGCCG | Homologous fragment |
| pJQ-Kan-nopM-down-R | GCGGCCGCTCTAGAAcTAGTGTTTCCAAATCCCCCTCGAGC | Homologous fragment |
| Kan-F | GCGTATTTCGTCTCGCTCA | Validation of mutation |
| Kan-R | GAGCGAGACGAAATACGCG | Validation of mutation |
| ttsI-up | gCTATCACAggTgTTTgggT | Validation of mutation |
| ttsI-down | GTGCGATCATTAGCGGATGA | Validation of mutation |
| nopA-up | ttagcgtttttttgtcctggagc | Validation of mutation |
| nopA-down | cggcgctgatcttaatgtt | Validation of mutation |
| nopJ-up | gAATTTTTTTgAAgAgCCAgATCg | Validation of mutation |
| nopJ-down | ggCgACAAAggTgTAAAAgTT | Validation of mutation |
| nopL-up | ACgTTCgAgCggACATAAg | Validation of mutation |
| nopL-down | TgACggATTCgCCCAAT | Validation of mutation |
| nopP-up | TAgCggCAgAAAgCTTTTTg | Validation of mutation |
| nopP-down | CAgAATATCgCCTgCATTCTCAA | Validation of mutation |
| nopZ-up | CTATgCAgAgTgCgAgTTTC | Validation of mutation |
| nopZ-down | gTTgCgTgACCTCTggT | Validation of mutation |
| nopM-up | CAGCTAAGCCTTGTAGAAGATTCG | Validation of mutation |
| nopM-down | GGGTCAGCCGGTTCT | Validation of mutation |
| pUB-Hyg-NopM-F | tgttgttgatgtgattacagtctagaATGAATGTACAA | Stable transformation |
| pUB-Hyg-NopM-R | tggtccttatagtcggtaccCAGCTCAAGACC | Stable transformation |
| pHyg-proNIN:NopM-F | ccaagctgggctgcagTTACACGTGGACGCAGC | Stable transformation |
| pHyg-proNIN:NopM-R | GGCCGTTGTACATTCATtctagaGCTAGCTGATCCAATTAAG | Stable transformation |
| NopM-600F | AGGTTGTCGCGGATTG | Stable transformation |
| NopM-1200R | GGTAACGTGAGAGACGTTCAAGAA | Stable transformation |
| p5XX-NopMC338A-cluc-F | agaacacgggggaCTCTAGAATGAATGTACAACGGCCC | Split-LUC assay |
| p5XX-NopMC338A-cluc-R | CCGCTGTTATCGGTACCCAGCTCAAGACCGCG | Split-LUC assay |
| p5XX-LjNFR5-cluc-F | agaacacgggggactCTAGAatggctgtcttctttcttacc | Split-LUC assay |
| p5XX-LjNFR5-cluc-R | gctcCCCGCTGTTATCGGTACCacgtgcagtaatggaagtcac | Split-LUC assay |
| p5XX-LjNFR1-cluc-F | agaacacgggggactCTAGAATGAAGCTAAAAACTGGTCTACTTTTG | Split-LUC assay |
| p5XX-LjNFR1-cluc-R | CCGCTGTTATCGGTACCTCTTCTCACAGACAGTAAATTTATGAGAGT | Split-LUC assay |
| p5XX-AtFLS2-cluc-F | agaacacgggggactctagaATGAAGTTACTCTCAAAGACCTTTTT | Split-LUC assay |
| p5XX-AtFLS2-cluc-R | cCCCGCTGTTATCGGTACCAACTTCTCGATCCTCGTTACG | Split-LUC assay |
| p5XX-NopMC338A-nluc-F | agaacacgggggaCTCTAGAATGAATGTACAACGGCCC | Split-LUC assay |
| p5XX-NopMC338A-nluc-R | CCCGCTGTTATCGGTACCCAGCTCAAGACCGCGA | Split-LUC assay |
| p5XX-LjNFR1K351E-cluc-F | agaacacgggggactCTAGAATGAAGCTAAAAACTGGTCTACTTTTG | Split-LUC assay |
| p5XX-LjNFR1K351E-cluc-R | CCGCTGTTATCGGTACCTCTTCTCACAGACAGTAAATTTATGAGAG | Split-LUC assay |
| pSPYCE-NopMC338A-F | agaacacgggggacTCTAGAATGAATGTACAACGGCCC | BIFC assay |
| pSPYCE-NopMC338A-R | GTCGACAGTACTATCGATGGATCCCAGCTCAAGACCGCG | BIFC assay |
| pSPYCE-LjNFR5-F | agaacacgggggacTCTAGAatggctgtcttctttcttacct | BIFC assay |
| pSPYCE-LjNFR5-R | GTCGACAGTACTATCGATGGATCCacgtgcagtaatggaagtca | BIFC assay |
| pSPYCE-LjNFR1-F | agaacacgggggactCTAGAATGAAGCTAAAAACTGGTCTACTTTTG | BIFC assay |
| pSPYCE-LjNFR1-R | GTCGACAGTACTATCGATGGATCCTCTTCTCACAGACAGTAAATTTA | BIFC assay |
| pSPYCE-AtFLS2-F | agaacacgggggactctagaATGAAGTTACTCTCAAAGACCTTTTT | BIFC assay |
| pSPYCE-AtFLS2-R | GTCGACAGTACTATCGATGGATCCAACTTCTCGATCCTCGT | BIFC assay |
| pSPYNE-NopMC338A-F | agaacacgggggactctagaATGAATGTACAACGGCCC | BIFC assay |
| pSPYNE-NopMC338A-R | gtcgacagtactatcgatggatccCAGCTCAAGACCGCG | BIFC assay |
| p5XX-NopMC338A-FLAG-F | agaacacgggggactCTAGAATGAATGTACAACGGCCC | Co-IP assay |
| p5XX-NopMC338A-FLAG-R | CCGCTGTTATCGGTACCCAGCTCAAGACCGCG | Co-IP assay |
| p5XX-LjNFR5-GFP-F | agaacacgggggactCTAGAatggctgtcttctttcttacc | Co-IP assay |
| p5XX-LjNFR5-GFP-R | tcccgggagcgGTACCacgtgcagtaatggaagtca | Co-IP assay |
| p5XX-LjNFR1-GFP-F | agaacacgggggactCTAGAATGAAGCTAAAAACTGGTCTACTTTTG | Co-IP assay |
| p5XX-LjNFR1-GFP-R | tcccgggagcgGTACCTCTTCTCACAGACAGTAAATTTATGAGAG | Co-IP assay |
| pMAL-Myc-F | AAGGATTTCAGAATTCGGATCCACCGATAACAGCGGGTTAATTAA | Pull down assay |
| pMAL-Myc-R | CGACGGCCAGTGCCAAGCTTcgatcggggaaattcgAGCTC | Pull down assay |
| pMAL-LjNFR5-Myc-F | GGAAGGATTTCAGAATTCGGATCCatggctgtcttctttcttacc | Pull down assay |
| pMAL-LjNFR5-Myc-R | CGACGGCCAGTGCCAAGCTTcaagtcttcctcggagattagct | Pull down assay |
| pACYduet-NopM-F | AATTCGAGCTCGGCGCGCATGAATGTACAACGGCCC | Pull down assay |
|  |  | Ubiquitination assay |
| pACYduet-NopM-R | TGTTCGACTTAAGCATTATGCGGCCGCgcgGTACCCAGCTCAA | Pull down assay |
| pCDFduet-LjNFR1-F | ACCACAGCCAGGATCCGAGATACCAGAAGAAGGAAGAAGAGAAAG | Ubiquitination assay |
| pCDFduet-LjNFR1-R | GGCGCGCCGAGCTCGcgatcggggaaattcgAGCT | Ubiquitination assay |
| pCDFduet-LjNFR5-F | ACCACAGCCAGGATCCGAATtatgtatactgccgcagaaagaagg | Ubiquitination assay |
| pCDFduet-LjNFR5-R | GGCGCGCCGAGCTCGAATTCcgatcggggaaattcgAGCTCt | Ubiquitination assay |
| pCDFduet-GmNFR5-F | GCCAGGATCCGAATTATTGTCTGAAAATGAAGACTTTGAATAGGAG | Ubiquitination assay |
| pCDFduet-GmNFR5-R | GGCGCGCCGAGCTCGAATTCcgatcggggaaa | Ubiquitination assay |
| pUB-NopM-FLAG-RFP-F | atctgttgttgatgtgattacagtctagaATGAATGTACAACGGCCC | Hairy root transformation |
| pUB-NopM-FLAG-RFP-R | cgatcgatggcgcgcctaggtacccgatcggggaaattcgAG | Hairy root transformation |
| NopM700-R | CATGCATGGCTGTCACCA | Validation of transformed plants |
| LjNINpro-F | CTTGATATTCATGCCCTCACTTTC | Validation of transformed plants |
| LjNFR5-800-F | tacgaaggatttcagcgatgag | Validation of transformed plants |
| LjNFR5-1200-R | gtatgttcatgcatgtattgcag | Validation of transformed plants |
| LjATPase-qRT-F | CAATGTCGCCAAGGCCCATGGTG | Gene expression analysis |
| LjATPase-qRT-R | AACACCACTCTCGATCATTTCTCTG | Gene expression analysis |
| NopM-qRT-F | TGGGAAAGAGTCATGAGGCG | Gene expression analysis |
| NopM-qRT-R | TGGGAAAGAGTCATGAGGCG | Gene Expression analysis |

**Table S2** **Mutants of S. fredii NGR234 Constructed in This Study.**

| Mutant^1^ | Characteristics^2^ |
| --- | --- |
| *ΩttsI* | NGR234 carrying the constructed ΩKm interposon (*pro:Kan*) at nucleotide position 15 of *TtsI* (Rif^r^, Km^r^). |
| *ΩnopA* | NGR234 carrying the ΩKm interposon at nucleotide position 24 of *NopA* (Rif^r^, Km^r^). |
| *ΩnopM* | NGR234 carrying the ΩKm interposon at nucleotide position 18 of *NopM* (Rif^r^, Km^r^). |
| Δ*nopTΩnopM* | Δ*nopT* carrying the ΩKm interposon at nucleotide position 18 of *NopM* (Rif^r^, Sp^r^, Km^r^). |
| Δ*nopTΩnopJ* | Δ*nopT* carrying the ΩKm interposon at nucleotide position 24 of *NopM* (Rif^r^, Sp^r^, Km^r^). |
| Δ*nopTΩnopL* | Δ*nopT* carrying the ΩKm interposon at nucleotide position 18 of *NopM* (Rif^r^, Sp^r^, Km^r^). |
| *ΩnopP* | NGR234 carrying the ΩKm interposon at nucleotide position 18 of *NopM* (Rif^r^, Km^r^). |
| Δ*nopTΩnopP* | Δ*nopT* carrying the ΩKm interposon at nucleotide position 18 of *NopM* (Rif^r^, Sp^r^, Km^r^). |
| *ΩnopZ* | NGR234 carrying the ΩKm interposon at nucleotide position 24 of *NopZ* (y4yJ) (Rif^r^, Km^r^). |
| Δ*nopTΩnopZ* | Δ*nopT* carrying the ΩKm interposon at nucleotide position 24 of *NopM* (Rif^r^, Sp^r^, Km^r^). |

^1^ In addition to indicated mutants, strains were GFP labeled using a derivative of pHC60 (FJ151627.1).

^2^ Rif^r^, resistance against rifampin; Km^r^, resistance against kanamycin; Sp^r^, resistance against spectinomycin.

**Table S3 LjNFR5 In Vitro Ubiquitination Sites Identified by Mass Spectrometry.**

Ubiquitination sites in LjNFR5 identified by mass spectrometry. Ubiquitinated lysine (K) residues are highlighted in red.

| Site | Peptide | Peptide hits |
| --- | --- | --- |
| K289 | TASSAETADKLLSGVSGYVSKPNVYEIDEIMEATK | 1 |
| K300 | LLSGVSGYVSKPNVYEIDEIMEATK | 8 |
| K314 | LLSGVSGYVSKPNVYEIDEIMEATK | 7 |
| K321 | DFSDECK | 1 |
| K328 | VGESVYKANIEGR | 12 |
| K342 | KIKEGGANEELK | 2 |
| K351 | EGGANEELKILQK | 6 |
| K355 | ILQKVNHGNLVK | 4 |
| K446 | DITTSNILLDSNFK | 4 |
| K488 | AMTTKENGEVVMLWK | 2 |
